# The glucocorticoid receptor is a critical regulator of muscle satellite cell quiescence

**DOI:** 10.1101/2023.08.27.555012

**Authors:** Rashida Rajgara, Hamood AlSudais, Aisha Saleh, Alex Brown, Ines Barrakad, Alexandre Blais, Nadine Wiper-Bergeron

## Abstract

Glucocorticoids are powerful anti-inflammatory medications that are associated with muscle atrophy. The effect of glucocorticoids in myofibers is well-studied, yet the role of the glucocorticoid receptor (GR), the primary mediator of glucocorticoid transcriptional responses, and the impact of glucocorticoid signalling in muscle stem cells (MuSCs), the adult progenitors responsible for regeneration, remain unknown. We developed a conditional null mouse model to knock out glucocorticoid receptor (GR) expression in MuSCs (GR^MuSC-/-^) and established that while GR is dispensable for muscle regeneration, it is a critical regulator of MuSC homeostasis. Loss of GR significantly increased cycling MuSCs as compared to controls in injury-naïve mice and on single EDL myofiber cultures, and as such, loss of GR in MuSCs leads to precocious activation and subsequent proliferation as compared to controls. Bulk RNA-sequencing from *in situ* fixed MuSCs from injury-naïve GR^MuSC-/-^ muscle identified a gene signature consistent with cells that have exited quiescence and undergone activation, with evidence of sexual dimorphism. Using ATAC-seq and footprinting we identify putative GR targets that promote quiescence. Thus, we advance the GR as a previously unrecognized crucial transcriptional regulator of gene expression in MuSCs whose activity is highest in quiescent cells and is essential to maintain that state.

## Introduction

Skeletal muscle regeneration following injury is assured by the presence of muscle-resident stem cells called satellite cell (MuSCs) (1, 2). Identified by their expression of Paired Box 7 (PAX7) and their localization to a niche between the sarcolemma and the basal lamina of muscle fibers, MuSCs are normally mitotically quiescent in healthy muscle. Quiescent MuSCs are characterized by decreased cell size and reduced histone methylation, RNA content, protein synthesis and metabolism (3). MuSCs are found in a highly specified niche, which consists of the adjacent myofiber, surrounding cells (fibro/adipogenic precursors, immune cells), blood vessels and the extracellular matrix (ECM), all of which interact with quiescent cells using indirect (various soluble factors) or direct (cell-cell; cell-matrix) contact (4). The dynamic interactions between MuSCs and their niche regulates their stem cell function.

Upon injury or muscle degeneration, MuSCs re-enter the cell cycle (5) and induce the expression of the myogenic regulatory factors MYOD1 and MYF5 (6). With terminal differentiation into myocytes, PAX7 is downregulated, the MRFs myogenin and MYF6 are expressed, and cells withdraw from the cell cycle irreversibly (5, 7, 8). Myocytes then fuse to form a multinucleated syncytium that becomes the myofiber or with existing fibers. Some myogenic precursor cells can escape differentiation, downregulate MYOD expression and self-renew, re-entering quiescence to maintain the MuSC population and ensure future regenerative capacity (5, 9, 10).

Glucocorticoids (GCs) are naturally occurring steroid hormones produced and released by the adrenal cortex in response to biological cues and stress such as fasting, exercise and disease (11). GCs regulate protein and glucose metabolism, development, immune function, skeletal growth, reproduction, cognition, and cardiovascular function (12). Synthetic GCs, by virtue of their powerful anti-inflammatory and immunosuppressive activity, are used therapeutically for numerous conditions (13). In 2020 alone, ∼20 million total prescriptions for prednisone, one of many synthetic glucocorticoids used therapeutically, were given to ∼9 million patients in North America, of which ∼3 million patients are chronic users (14). Despite their therapeutic potential, the use of GCs has serious side effects including muscle wasting, which affects up to 60% of patients (15). Indeed, high circulating GCs decrease skeletal muscle protein anabolism and increase protein catabolism to produce free amino acids, a substrate for gluconeogenesis by the liver. Prolonged elevated glucocorticoid levels due to pathology (Cushing’s syndrome, cancer cachexia or sepsis) or exogenous use results in loss of muscle mass and muscle weakness (15, 16).

Glucocorticoid action is mediated by the glucocorticoid receptor (encoded by the *Nr3c1* gene), which is a member of the nuclear hormone receptor superfamily of ligand-activated transcription factors. Upon ligand binding, the GR translocates from the cytoplasm to the nucleus where it interacts with GC response elements in the regulatory regions of target genes (17) directly or via interaction with sequence-specific transcription factors to regulate transcription independently of direct DNA-binding (18, 19). In muscle, the GR and MYOD1 interact with the CpG-bound transcription factor NFR1 to activate gene expression of the skeletal muscle program (20) and the GR negatively regulates hypertrophy and striated muscle contraction in mice by downregulating anabolic pathways (20). While a role for the GR in the myofiber has been described, its importance in MuSC function has not been thoroughly investigated (20). Herein, we report the phenotype of a novel mouse model in which the GR is knocked out in MuSCs and establish that the GR is necessary for MuSC quiescence.

## Materials and Methods

### Biological Resources

#### Generation of GR conditional knockout mice and animal care

*Nr3c1*^fl/fl^ floxed mice (21) (B6.Cg-Nr3c1tm1.1Jda/J, The Jackson laboratory, <u>stock # 021021</u>) were crossed with *Pax7*^CreER^ mice (B6.Cg-Pax7tm1(cre/ERT2)Gaka/J, The Jackson laboratory, stock <u>#017763</u>) (22) to produce *Nr3c1*^+/fl^*;Pax7*^creER/+^ and *Nr3c1*^+/fl^*;Pax7*^+/+^ progeny. The *Nr3c1*^+/fl^*; Pax7*^CreER/+^ male studs were then crossed with *Nr3c1*^fl/fl^*; Pax7*^+/+^ females to produce WT (*Nr3c1*^fl/fl^*; Pax7*^+/+)^ and GR^MuSC-/-^ (*Nr3c1*^fl/fl^*; Pax7*^CreER/+^) pups at Mendelian ratios. Temporal control of the GR knockout was achieved via 5 daily intraperitoneal (i.p.) injection of tamoxifen (1.5-2 mg/20 g body weight) at 4-6 weeks of age. Mice were maintained at 22°C with 30% relative humidity on a 12-hr light/dark cycle and provided food and water ad libitum. Animal experimentation performed at the University of Ottawa was approved by the University of Ottawa Animal Care Committee and conducted in accordance with the guidelines set out by the Canadian Council on Animal Care. For ATAC-seq, mice were kept in accordance with IGBMC guidelines for the care and use of laboratory animals and in accordance with National Animal Care Guidelines (European Commission directive 86/609/CEE; French decree no.87–848). All procedures were approved by the French national ethics committee.

#### Isolation and culture of primary myoblasts

Primary myoblasts from adult (aged 6-8 weeks) WT and GR^MuSC-/-^ conditional knockout mice hindlimb muscles were isolated by magnetic activated cell sorting (MACS) as previously described (23). Cells were washed with serum-free media and enriched for myoblasts by selective plating to eliminate contaminating fibroblasts for 30-45 min. Primary myoblasts were cultured on matrigel-coated plates in growth media (GM; Dulbecco’s modified Eagle’s medium [DMEM] containing 20% fetal bovine serum [FBS], 10% horse serum [HS]) supplemented with basic fibroblast growth factor (10 ng/ml) and hepatocyte growth factor (2 ng/ml). *In vitro* excision of Nr3c1 exon 3 was achieved by treating primary myoblasts with 4-OH-tamoxifen (Sigma Aldrich, H7904).

### Cardiotoxin injury and analysis of muscle histology

Seven-week-old mice were anesthetized with isofluorane, and 30 μL of 10 μM CTX (Latoxan, France) in PBS was injected intramuscularly into the left Tibialis anterior (TA) muscle, the right TA was left uninjured. Mice were sacrificed by cervical dislocation, 7- and 28-days post injury and the dissected muscles were flash frozen using Optimal Cutting Temperature compound (OCT) dipped in isopentane cooled by liquid nitrogen. Eight μm-thick sections of TA muscle were fixed on charged slides using the HM525NX cryostat and stained with hematoxylin & eosin following fixation in 4% paraformaldehyde (PFA) for 15 min (Louise Pelletier Histology Core Facility, University of Ottawa). Images were acquired on the EVOS FLAuto2 (Cell Biology & Image Acquisition Core Facility, University of Ottawa). A minimum of 500 fibers were counted for analysis of the average fiber cross-sectional area using FIJI (Fiji Is Just ImageJ) for quantification.

### Immunofluorescence

Single EDL myofibers were fixed in 4% paraformaldehyde (PFA) in PBS and 1% glycine. Fibers were blocked in PBS containing 0.2% Triton X-100, 2% BSA, 5% normal donkey serum and 1% azide. TA sections (cryopreserved at −80°C) were dehydrated at 65°C on a heating block and fixed in 4% paraformaldehyde (PFA). Membrane permeabilization was carried out using 0.5% Triton X-100 in PBS and blocked using 0.1% TritonX-100 in PBS with 5% normal donkey serum (Cedarlane, 017-000-121). Specifically, for PAX7 staining, antigen retrieval was performed in a citrate-buffered solution (pH 6.0) at 92°C for 30 min prior to permeabilization. Next, the sections were blocked for 30-1hr, following primary antibody incubation at 4°C overnight. Primary myoblasts were fixed using PFA and permeabilized in PBS containing 0.5% Triton X-100 and 10% normal donkey serum (Cedarlane, 017-000-121). Detection was performed according to standard procedures using the following primary antibodies: PAX7-c (DSHB), Glucocorticoid receptor (Cell Signalling, D6H2L), MF-20 (DSHB), Dystrophin (Abcam, ab15277), Laminin (Abcam, ab11575), Ki67 (Abcam, ab15580), and BrdU (Shp pAb to Biotin) (Abcam, ab2284). Detection of immunoconjugates was achieved using, (1) biotinylated secondaries such as donkey anti-mouse IGG (H+L) Fab fragments (Jackson ImmunoResearch Laboratories, 715-066-150) with Cy3-streptavidin (Jackson ImmunoResearch Laboratories, 016-160-084), or Alexa Fluor 488-Streptavidin (Jackson ImmunoResearch Laboratories, S11223) for biotinylated BrdU and (2) non-biotinylated secondary antibodies such as Dylight 488 donkey anti-rabbit (Jackson ImmunoResearch Laboratories, 016-540-084) and Cy3 anti-mouse (Jackson ImmunoResearch Laboratories, 715-166-150). Nuclei were counterstained with DAPI and mounted in vectashield antifade mounting medium (Vectorlabs, H-1000). Images were taken on the Zeiss AxioImager.M2 Microscope (Cell Biology & Image Acquisition Core Facility).

### *In vivo* BrdU labeling

Six-week-old mice received 5 daily i.p. injections of tamoxifen to knockout the GR. Two weeks following the last injection, *Nr3c1*^fl/fl^; *Pax7*^+/CreER^ (GR^MuSC-/-^) and *Nr3c1*^fl/fl^; *Pax7*^+/+^ (WT) male and female mice were intraperitoneally injected once with 100 mg/kg of body weight of BrdU and TA muscles and hindlimbs were collected 24 hrs later.

### ATAC-seq

MuSCs were isolated and fixed from C57BL/6 mice (8-10 weeks of age) as described in Machado *et al*. (24). In the case of cells that were processed fresh (non-fixed), certain samples were aliquoted in two where one aliquot was left untouched while the other was cross-linked after FACS (thus after *in vitro* activation) using 1% formaldehyde for 5 minutes at room temperature, followed by quenching with glycine (0.125 M) for 5 minutes at room temperature and rinsing twice with PBS. Cells that were grown *in vitro* were plated on Matrigel-coated culture dishes in growth medium (GM; Dulbecco’s modified Eagle’s medium [DMEM] containing 20% fetal bovine serum [FBS], 10% horse serum [HS]) and harvested 24 hours later.

ATAC-seq was performed using the OMNI-ATAC variant of the protocol (25). After tagmentation, cross-links were reversed as previously described (26) by adjusting the samples to 66 mM Tris pH 8, 1.3 % (v/v) SDS and 266 mM NaCl and heating samples at 65°C for at least 8 hours. PCR amplification of the libraries was performed using NEBnext Q5 Ultra II master mix (NEB) and the last sample cleanup was performed by double-sided PEG-magnetic bead purification to eliminate PCR primers and fragments above 1 kb in size. Paired-end sequencing was performed on HiSeq4000 instruments at the GenomEast genomics platform at the Institut de Génétique et de Biologie Moléculaire et Cellulaire (IGBMC, Illkirch-Graffenstaden, France). Raw sequencing reads have been deposited to the National Center for Biotechnology Information (NCBI) Gene expression Omnibus (GEO) under accession number GSE234936.

### ATAC-seq data processing and analysis

Raw sequencing files were processed as described in Madden *et al* (27). Alignment was performed on version mm9 of the mouse genome. Differential accessibility analysis was performed on the union of peaks called by MACS2 (version 2.2.7.1) (27) on each sample. Tnp (Tn5 transposase) insertion sites were quantified at each peak using the featureCounts function of Subread (version 1.34.7) (28, 29) specifying to count each mate of the read pairs independently, and to count only the 5’-most nucleotide. R (version 4.2.1) and Bioconductor (version 3.15) with the edgeR package (version 3.38.4) (30) were used to identify differentially accessible regions (DARs). TMM adjustment of library sizes was performed and followed by applying RUVseq (version 1.30.0) to remove unwanted variation (31). Empirical testing and evaluation by PCA plots and p-value distribution histograms (32) showed satisfactory performance using the RUVr algorithm with a k value of 2. Heatmaps were generated using R/Bioconductor and the ComplexHeatmap package (version 2.12.1) (33).

Transcription factor footprinting was performed using RGT-HINT (version 1.0.1) and the HINT-ATAC function (34). The TF motif database contained 1297 motifs from the HOCOMOCO v10 database (35) and 1099 motifs discovered de novo by MEME-ChIP (version 5.5.0) (36) as enriched in the ATAC-seq signal peaks unique to one cell state (e.g., quiescent) compared to the other two (e.g., in vitro activated or grown). Footprinting was limited to ATAC-seq signal peaks. A total of 2395 TFs had a motif match among footprint regions, and this list was reduced to 2020 by removing TFs with less than 1500 footprints or more than 15000, genome wide. The change in activity of all TFs between two cell states was calculated, normality of their distribution was verified using the Shapiro-Wilk test (37) and p-values were calculated. TFs with a p-value smaller than 0.05 were noted as having significantly different activity.

To align this study’s ATAC-seq data with mRNA expression levels from comparable samples, RNA-seq data from Machado *et al*. was downloaded from the NCBI Gene Expression Omnibus accession GSE103164 using the provided count matrix and processed using a standard pipeline (27). Unwanted variation was reduced using the RUVr algorithm and a k value of 2. Normalized counts and log2 fold-change values were recorded to be analyzed alongside the ATAC-seq data. DA regions were annotated to genes using the ChIPpeakAnno R/Bioconductor package (version 3.30.1) (38), assigning the nearest gene transcriptional start site within a maximum distance of 50 kb as the one “target” gene. When multiple DA regions were annotated to the same gene, all regions were retained such that the same gene can appear multiple times in the aggregate results.

### Single EDL myofiber isolation

Myofibers were isolated from the Extensor digitorum longus (EDL) muscle as described previously (39). Isolated myofibers were fixed and stained immediately upon isolation (∼2h post dissociation) or cultured for 3 days (∼72h) in suspension media before immunostaining to quantify myogenic cell populations.

### Western analysis

Total protein was collected from WT and GR^MuSC-/-^ primary myoblasts by homogenization in IPH lysis buffer (50 mM Tris–HCl pH 7.5, 150 mM NaCl, 5 mM EDTA, 0.5% NP-40) supplemented with 100 mM PMSF and 100 mM DTT (Dithiothreitol, Sigma Aldrich). Protein samples were resolved on an 12% SDS-polyacrylamide gel electrophoresis and transferred to polyvinylidene fluoride membrane for detection with the following antibodies: anti-Glucocorticoid Receptor D6H2L (1:1000 dilution, Cell Signaling, New England BioLabs #12041), anti-PAX7c (1:500 dilution, Developmental Studies Hybridoma Bank, DSHB), anti-MyoD5.8A (1:200 dilution, SantaCruz, sc-32758) and anti-Cyclophilin B (1:1000 dilution, Abcam, ab16045) antibodies. Glucocorticoid receptor (GR), PAX7, MYOD and Cyclophilin B were visualized using horseradish peroxide (HRP)-conjugated anti-rabbit (for GR and Cyclophilin B) and anti-mouse for (PAX7 and MYOD) at a 1:5000 dilution and detected by chemiluminescence.

### Isolation of in situ fixed muscle satellite cells

Muscle stem cells from fixed tissues were isolated according to Machado *et al.* (24) using the digestion protocol developed by Baker *et al* (40). Fixed MuSCs from WT and GR^MuSC-/-^ mice were purified by FACS via gating for Streptavidin BV421-positive muscle stem cells conjugated to the positive marker, (VCAM-α mCD106 receptor) present on these cells, using a 4-laser Sony SH-800 cell sorter.

Antibody Panel used for FACS isolation of MuSCs from in situ fixed muscle:

| Selection | Antibody | Antibody conjugate | Catalogue number | Concentration |
| --- | --- | --- | --- | --- |
| Negative | CD31 | PE-Cy7 | 561410 (BD Biosciences) | 0.2 mg/mL |
|  | CD45 | PE-Cy7 | 552848 (BD Bioscience) | 0.2 mg/mL |
|  | Sca1 | PE | 553108 (BD Bioscience) | 0.2 mg/mL |
| Positive | VCAM (a-Mcd106) | Biotin | 31-0057-05 (AbLab) | 1.0 mg/mL |
|  |  | Streptavidin BV421 | 563259 (BD Bioscience) | 0.1 mg/mL |

### RNA-seq analysis

The RNA from in situ fixed and sorted cells (each replicate derived from a minimum of 3 mice of the same sex and age) was isolated using the RecoverAll™ Total Nucleic Acid Isolation Kit for formalin-fixed parafilm embedded (FFPE) samples (Invitrogen, cat#AM1975). Library preparation, PCR amplification and sequencing was performed at the Donnelly Sequencing Center at the University of Toronto. Raw reads have been deposited to the NCBI GEO under accession number GSE235039.

FASTQ files from RNA-seq were processed using a standard pipeline. Briefly, reads were processed first with UMI-tools (v. 1.1.2) (41) to extract unique molecular identifiers (UMIs). Then fastp (v. 0.20.1) (42) was used to trim adapters and low-quality sections. Reads were aligned to the mouse genome (mm10 assembly) using STAR (v. 2.7.8a) (43) and an index was created using a gene model GTF file (Mus_musculus.GRCm38.102.gtf, from ENSEMBL). UMI information was used to remove reads marked as duplicates. Reads overlapping exons were counted using Subread featureCounts (v. 2.0.1) (29) and counts were summarized over genes. Read counts were analyzed in R/Bioconductor using edgeR (v. 3.38.4) (30) employing the TMM method to adjust library sizes. Genes with transcript shorter than 300 nucleotides or with expression less than three-fold from the median in at least 2 samples were removed from the analysis. Unwanted variation from the count table was removed using RUVseq (v. 1.30.0) (31). Empirical tests using the RUVr, RUVs and RUVg algorithms and different values of k showed good performance with all methods and RUVs with k=2 was chosen. Gene annotation term over-representation analyses and gene set enrichment analyses were performed in R/Bioconductor using the clusterProfiler package (v. 4.4.4) (44). Heatmaps were generated using ComplexHeatmaps (v. 2.12.1) (45).

### Statistical analysis

All analysis was performed using a blinded design. Statistical differences between two means were calculated using the two-tailed Student’s t-test for a minimum of three biological repeats. For multiple means, one-way or two-way ANOVA was conducted. If significant differences were identified, multiple comparison analysis was performed using a Tukey’s post hoc test. Distributions were analyzed by Chi-square goodness of fit analysis. P-values of less than 0.05 were considered significant, *p<0.05, **p<0.01, ***p<0.001. GraphPad Prism 9.4.1 was used for statistical analysis.

### Data Availability/Sequence Data Resources

Sequencing datasets have been deposited on National Center for Biotechnology Information (NCBI) Gene expression Omnibus (GEO) using the following accession numbers:

ATAC-seq data: GSE234936

RNA-seq data: GSE235039

### Websites/Database Referencing

The RNA-seq of Machado et al. (in situ-fixed MuSCs) was downloaded from the NCBI GEO, https://www.ncbi.nlm.nih.gov/geo/query/acc.cgi?acc=GSE103164.

The microarray data of Rodgers et al. (G-alert genes in MuSCs) was downloaded from the NCBI GEO, https://www.ncbi.nlm.nih.gov/geo/query/acc.cgi?acc=GSE55490.

TF binding motifs were from the HOCOMOCO database version 10 and were downloaded as part of the RGT-HINT distribution.

They are available at https://hocomoco11.autosome.org/downloads_v10.

The mouse gene model was downloaded from the ENSEMBL at https://ftp.ensembl.org/pub/release-102/gtf/mus_musculus/Mus_musculus.GRCm38.102.chr.gtf.gz.

### Figure preparation

Photomicrographs from the results section were processed using FIJI (FIJI Is Just ImageJ) and the figures were assembled in Adobe Illustrator CS5.1™. The graphical abstract was created using Biorender.com.

## Results

### GR footprints are associated with genomic regions active in MuSC quiescence

To identify transcriptional regulators involved in the maintenance of mitotic quiescence of muscle satellite cells, we performed ATAC-seq on MuSCs isolated from C57BL/6 mice that were *in situ* fixed, freshly isolated or isolated and cultured for 2 hours according to the procedure described in Machado *et al* (24) (Fig. 1A). Overall, the samples clustered primarily by cell state (quiescence, early activation or growth) and fixation had a negligible effect on DNA accessibility profiles (Supplementary Fig. 1A, B). We identified regions of differential accessibility in quiescent versus early activated MuSCs including at genes known to be involved in MuSC-specific functions such as cell activation or myoblast differentiation (e.g., *Pax7*, *CalcR*, *FosB, Chrna1*) (Supplementary Figure 1C). Regions characterized by significant differential accessibility between *in situ*-fixed and *in vitro*-activated cells clustered best by k-means with k=4 (Fig. 1A, lefthand side). To correlate ATAC-seq peaks with gene expression, mRNA expression of the closest gene (within a 50 kb distance to the transcriptional start site) was identified using the Machado *et al*. dataset, which profiled gene expression in samples prepared under similar conditions to the ATAC-seq samples (24). Each ATAC-seq cluster was re-clustered based on mRNA levels using k-means and k=3 producing a heatmap showing ATAC-seq signal or RNA-seq signal in each of these twelve clusters (Fig. 1A, righthand side). Boxplots reveal the log2 fold-change in DNA accessibility (from ATAC-seq) or mRNA levels (from RNA-seq) between freshly isolated and quiescent cells for the regions and genes represented in each of the twelve clusters (see Supplemental Fig. 2). Interestingly, two clusters (9 and 12) showing increased DNA accessibility and mRNA levels during *in vitro* activation were enriched for gene ontology terms related to the response to GCs, cell growth and various terms related to muscle cell differentiation, suggesting that glucocorticoid action correlates with the regulation of cell growth and differentiation of MuSCs. A third cluster (cluster 2) featuring high accessibility and low gene expression in quiescent cells was similarly enriched for gene ontology terms related to muscle cell proliferation and glucocorticoid action (Fig. 1A).

**Figure 1.**
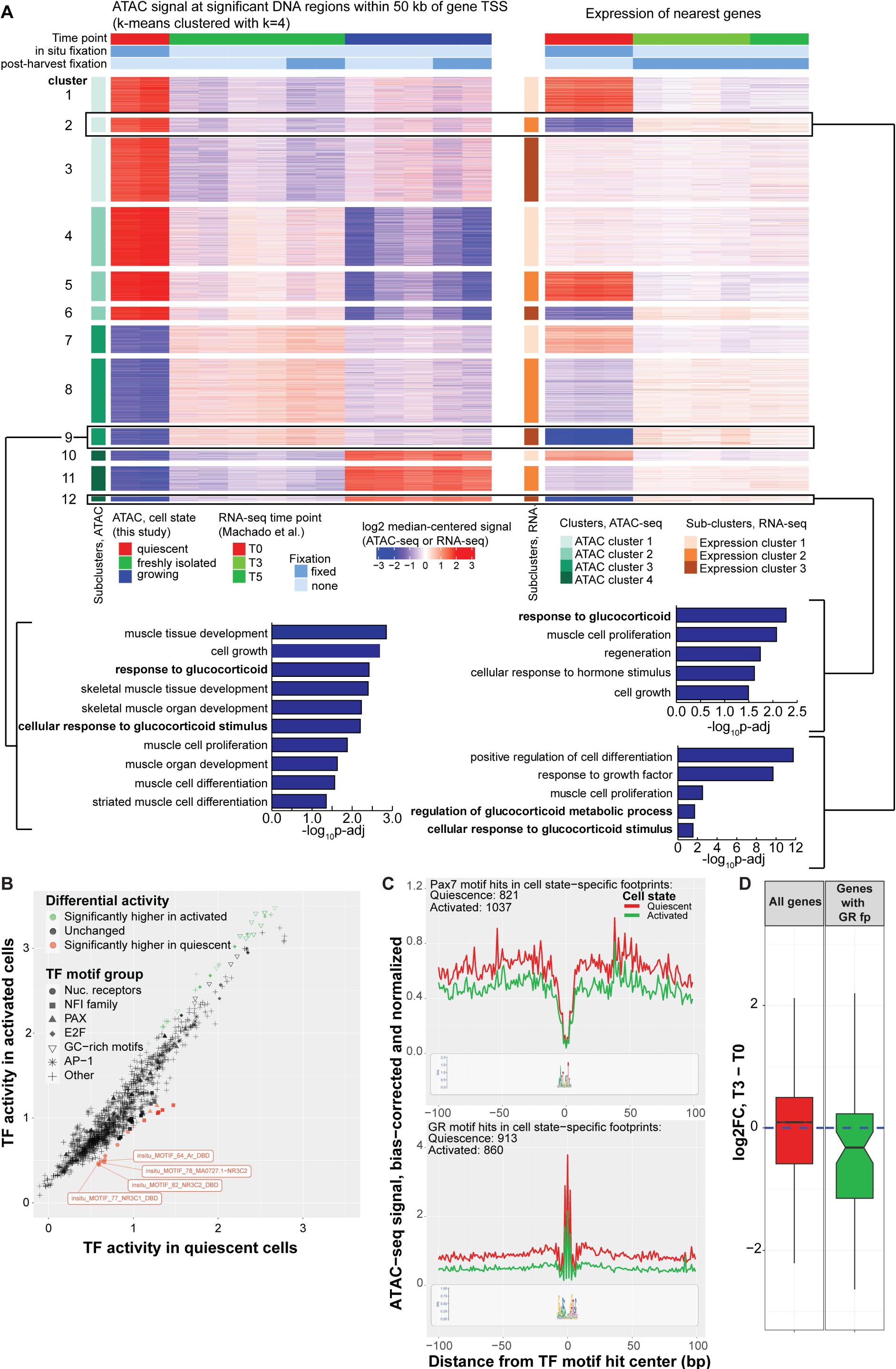
Transcription factor footprinting identifies GR as more active in quiescent MuSCs than early activated cells. **(A)** ATAC-seq signal at DNA regions in *in situ* fixed MuSCs (quiescent), freshly isolated and growing k-means clustered and compared to RNA-seq gene expression from Machado et al (GSE103162) where T0 is quiescent *in situ* fixed, T3 is freshly isolated and T5 is growing MuSCs. Analysis of three clusters (boxed) reveals enrichment for glucocorticoid signalling, cell growth/proliferation and muscle development. **(B)** Dot plot of transcription factor motifs identified in ATAC-seq using transcription factor footprinting where each dot represents a TF motif surveyed (green, more active in freshly sorted cells; orange, more active in quiescent cells). **(C)** Footprints for PAX7 and GR indicate higher activity in quiescent cells compared to early activation cells. Normalized and bias-corrected ATAC-seq signal in a 200 bp window centered on Pax7 or GR footprint sites in quiescent and early activation MuSCs. The consensus DNA sequences over the 200 bp regions are shown below. **(D)** Genes located near GR footprints are more downregulated in activated cells versus quiescent cells, compared to genes without GR footprints. Notches in boxplots indicate the 95% confidence interval.

To explore this, genomic regions with significant ATAC-seq peaks were assessed for the presence of transcription factor binding motifs (Fig. 1B). We identified 3 classes of transcription factors whose motifs were preferentially footprinted in quiescent MuSCs (*in situ* fixed) versus early activated cells: PAX3/7, NFI and the response element common to steroid hormone receptors including the GR (Fig. 1B). We found the glucocorticoid response element footprints significantly enriched in regions that are more active in quiescent cells in comparison to early activated cells (Fig. 1B, C). Indeed, using the Machado *et al* RNA-seq dataset to identify RNA transcripts present in quiescent and activated satellite cells, we find that as a group, genes in the vicinity of a GR footprint were found to be less expressed in activated MuSCs than in quiescent ones, a phenomenon not observed when all detected genes were considered. (Fig. 1D). These results suggest that the GR is a previously unrecognized important transcriptional regulator of gene expression in MuSCs and that its activity is highest in quiescent cells.

### Loss of GR expression does not negatively impact muscle regeneration following injury but results in fewer MuSCs and larger myofibers

To investigate a role for the GR in MuSC quiescence, activation and differentiation, we generated a temporally controlled conditional knockout of the GR in MuSCs using *Pax7*-driven CreER expression. Tamoxifen-mediated knockdown of the GR in *Nr3c1^fl/fl^*; *Pax7*^+/*CreER*^ mice (GR^MuSC-/-^) and their littermate controls (WT: *Nr3c1^fl/fl^; Pax7^+/+^*) was verified by western blot of MuSCs isolated from hindlimb muscle at experimental endpoint (Fig. 2A). We assessed muscle regeneration *in vivo* following a single cardiotoxin (CTX)-induced injury of the left tibialis anterior (TA) muscle one week following GR knockout. The right TA (contralateral, CLL) remained uninjured as a control. Seven days post-injury, both the GR^MuSC-/-^ and WT controls had elicited a strong regenerative response as evidenced by the presence of centrally located nuclei (Fig. 2B). The average myofiber cross-sectional area (XSA) of the right (CLL) TA muscles from WT and GR^MuSC-/-^ mice were equivalent (Fig. 2B, C) and following injury, both genotypes had regenerated to ∼55% of their respective uninjured controls (Fig. 2B, C). Thus, loss of GR expression does not negatively impact muscle regeneration following injury.

**Figure 2.**
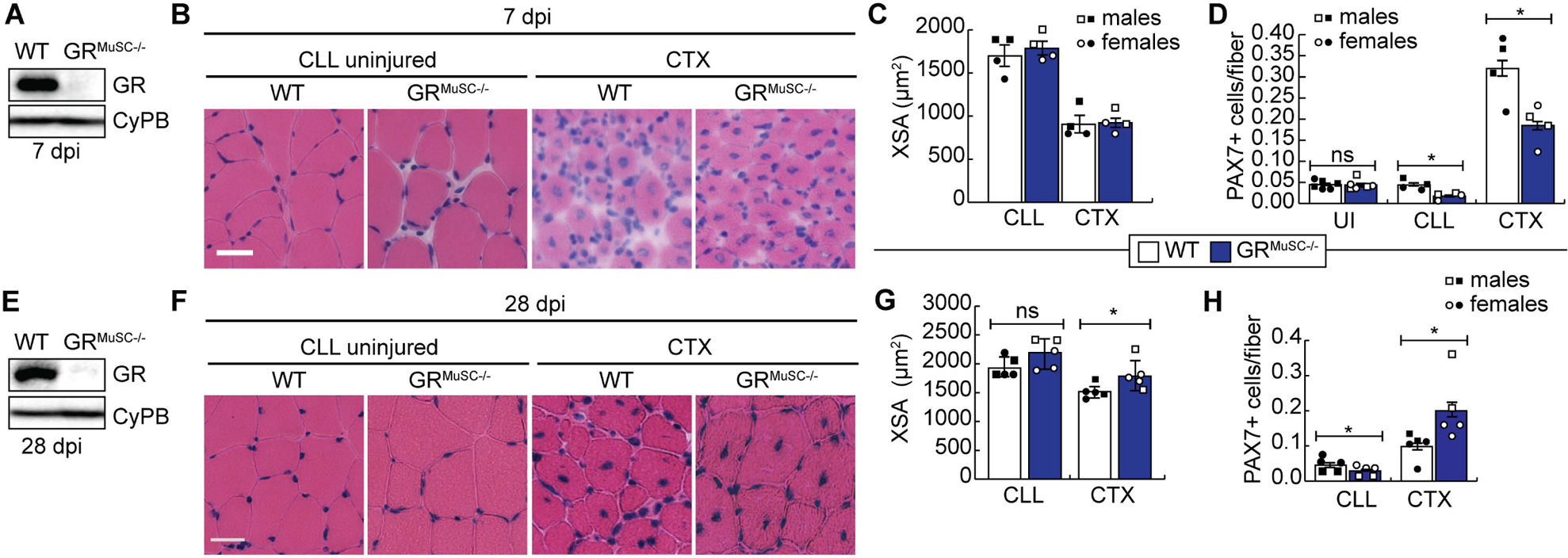
Loss of the GR in MuSCs results in PAX7+ cell number abnormalities and larger fibers in injured muscle. **(A)** Western blot of GR expression in primary myoblasts isolated from GR^MuSC-/-^ and WT hindlimb muscle 2 weeks after completion of tamoxifen administration. Cyclophilin B (CyPB) is a loading control. **(B)** H&E-stained TA muscle sections from WT and GR^MuSC-/-^ mice 7 days post cardiotoxin injury (7 dpi) compared to uninjured contralateral muscle (CLL). Scale bar = 50 μm**. (C)** Mean fiber cross-sectional area (XSA) from B. **(D)** Average number of PAX7+ MuSCs per fiber in uninjured TA (UI, injury-naïve controls), contralateral uninjured TA (CLL) and CTX-injured TA muscles from B. Data from male animals are indicated with square data points and female animals with circle data points. n=4 biological pairs, *p<0.05. Data is mean ± SEM. **(E)** Western blot of GR expression in primary myoblasts isolated from GR^MuSC-/-^ and WT hindlimb 28 days post-injury. Cyclophilin B (CyPB) is a loading control. **(F)** H&E stained cardiotoxin (CTX)-injured TA muscle sections from WT and GR^MuSC-/-^ mice at 28 dpi compared to CLL uninjured muscle. Scale bar = 50 μm. **(G)** Mean fiber cross-sectional area (XSA) of fibers from the right (CLL) and left (injured, CTX) TA muscle of mice treated as in F. **(H)** Average number of PAX7+ MuSCs per fiber in contralateral uninjured TA (CLL) and CTX-injured TA muscles from F. n=5 biological pairs, *p<0.05, two-tailed. Data is mean ± SEM.

Despite equivalent regenerative responses at 7 days post-injury, the PAX7^+^ cell numbers were significantly lower in injured GR^MuSC-/-^ muscle (∼40% reduction) (Fig. 2D). Further, we observed a significant 50% reduction in PAX7^+^ cells in the CLL TA muscle (Fig. 2D). Overall, MuSC numbers in the GR^MuSC-/-^ mouse expanded following injury (CTX vs CLL) by 9.5-fold, similar to controls (8-fold) suggesting no defects in cell proliferation (Fig. 2D). Given the dramatic reduction of PAX7^+^ cells in the contralateral TA only 2 weeks after GR expression knockout, we quantified PAX7^+^ cell numbers in injury-naïve mice (UI) at the same age and compared these to the cardiotoxin injured and uninjured contralateral limb (CLL). In injury-naïve muscle, we observed no significant changes in the number of PAX7^+^ MuSCs between WT and GR^MuSC-/-^ mice suggesting that injury provides a context that precipitates loss of PAX7^+^ cells in the absence of the GR (Fig. 2D).

To determine if the observed reduction in PAX7^+^ number impairs the regenerative process in the longer term, we assessed regeneration at 28 days after cardiotoxin injury (Fig. 2E-H). Knockout of the GR in MuSCs was verified by Western blot of isolated MuSCs from non-TA hindlimb muscle at experimental endpoint (Fig. 2E). Repair progressed in both the WT and GR^MuSC-/-^ muscle following injury and at 28 dpi the GR^MuSC-/-^ muscle had a significant increase in the average injured muscle fiber cross-sectional area as compared to controls (Fig. 2F, G). At 28dpi, in the uninjured contralateral limb (CLL), we found that GR^MuSC-/-^ mice had 33% fewer PAX7^+^ cells as compared to WT, as observed in early repair (Fig. 2H). In the injured TA, PAX7^+^ cell numbers were superior to controls due largely to decreasing MuSC numbers in the WT (Fig. 2H) at this time point.

### Single myofiber cultures reveal precocious differentiation and reduced PAX7^+^ stem cells with loss of GR

To study the role of the GR in the regulation of myogenic cell fate in resting muscle, EDL myofibers were collected from WT and GR^MuSC-/-^ mice two weeks following TAM treatment and GR knockout was confirmed by Western blot of MACS-isolated MuSCs from the remainder of the hindlimb muscles (Fig. 3A). EDL myofibers were immunostained for PAX7 and MYOD 2h post dissociation (Fig. 3B). We first quantified all myogenic cells (PAX7^+^ and/or MYOD^+^) and found a 2-fold increase on the GR^MuSC-/-^ myofibers as compared to WT (Fig. 3C) 2 hours post-isolation. At this time point, little MuSC activation was anticipated however, on GR^MuSC-/-^ myofibers, we observed a significant increase in PAX7+MYOD-cells, and while failing to meet statistical significance, larger populations of proliferating and differentiating cells were detected (Fig. 3D). We did, however, observe that while ∼80% of WT myogenic cells were PAX7^+^MYOD^-^ in GR^MuSC-/-^ muscle only ∼68% were quiescent (Fig. 3E). We also observed a 2-fold increase in proliferating cells at this time point with the overall distribution of cells, using the Chi-square goodness of fit, in the GR^MuSC-/-^ statistically different from WT (Fig. 3e). When fibers were stained for PAX7 and Ki67, a marker for cycling cells, the PAX7^+^Ki67^+^ population was ∼30% greater on GR^MuSC-/-^ myofibers as compared to WT controls (Fig. 3F, G), consistent with increased activation of MuSC cells. At 72 hours post-isolation, when MuSCs are expected to be activated and differentiating, we observed a similar increase in total myogenic cells per fiber in the GR^MuSC-/-^ with a particular increase in the PAX7^+^MYOD^+^ population (Fig. 3H-J). While no significant differences were observed in terms of number of cells per fiber for the PAX7^+^MYOD^-^ population (Fig.3J), the distribution of cells as PAX7^+^MYOD^-^, PAX7+MYOD+ and PAX7-MYOD+ was significantly different from WT, with a ∼25% decrease in PAX7+MYOD-population in GR^MuSC-/-^ mice 72 hours post-dissociation as compared to WT (Fig. 3K). Hence, our data suggests that loss of the GR leads to an increase in the percentage of Ki67^+^ cycling cells and PAX7^+^MYOD^+^ activated MuSCs population at the expense of the PAX7^+^MYOD^-^ population.

**Figure 3.**
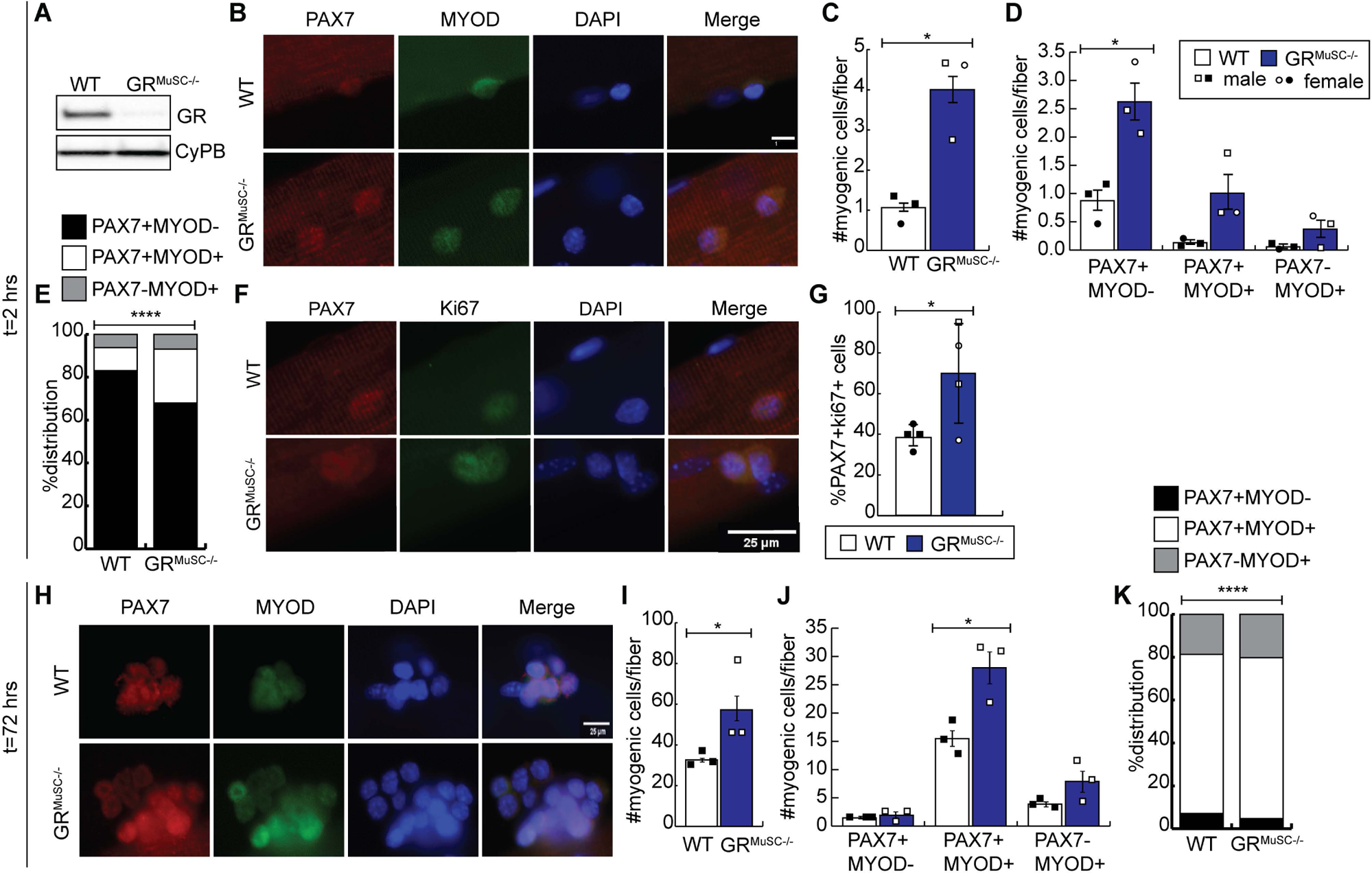
Loss of the GR leads to increased cell proliferation post-activation in ex vivo single EDL myofibers. GR^MuSC-/-^ and WT mice were treated with tamoxifen for 5 days and 2 weeks after the last injection, myofibers were isolated from the extensor digitorum longus (EDL) of each mouse and left in suspension for 2h or 72h before immunostaining. **(A)** Western analysis of GR and MYOD protein expression in MuSCs isolated from the non-EDL hind limb muscles of WT and GR^MuSC-/-^ mice at endpoint. **(B)** Representative images of myofibers stained for PAX7 (red) and MYOD (green) at t=2hrs. Nuclei were counterstained with DAPI (blue). Scale bar = 25 μm. **(C)** Quantification of total myogenic cells per fiber at 2h post-dissociation. n=3 biological pairs. **(D)** Number of quiescent/self-renewing (PAX7^+^MYOD^−^), activated/proliferating (PAX7^+^MYOD^+^) and differentiating (PAX7^−^MYOD^+^) myoblasts per myofiber at T2h from dissociation as in B. **(E)** Percent quiescent/self-renewing (PAX7^+^MYOD^−^), activated/proliferating (PAX7^+^MYOD^+^) and differentiating (PAX7^−^MYOD^+^) myoblasts on single myofibers at T2h. **(F)** Representative images of myofibers stained for PAX7 (red) and Ki67 (green) at t=72hrs. Nuclei were counterstained with DAPI (blue). Scale bar = 25 μm. **(G)** Quantification of PAX7+Ki67+ cells from F. n=4 **(H)** Representative images of myofibers stained for PAX7 (red) and MYOD (green) at t=72hrs. Nuclei were counterstained with DAPI (blue). Scale bar = 25 μm. **(I)** Quantification of total myogenic cells per fiber at 72h post-dissociation. **(J)** Number of quiescent/self-renewing (PAX7^+^MYOD^−^), activated/proliferating (PAX7^+^MYOD^+^) and differentiating (PAX7^−^MYOD^+^) myoblasts per myofiber at T72h. **(K)** Percent quiescent/self-renewing (PAX7^+^MYOD^−^), activated/proliferating (PAX7+MYOD+) and differentiating (PAX7^−^MYOD^+^) myoblasts on single myofibers at T72h from h. n= 3 biological pairs. Data is represented as mean ± SEM, *p < 0.05, two-tailed.

### MuSC from females and males re-enter the cell cycle differently

Given that loss of the GR in MuSCs leads to increased proliferation and an increase in cycling cells (Fig. 3), we postulated that the GR was required for the maintenance of MuSC quiescence. To understand how the GR regulates the MuSC transcriptome and discover novel targets during quiescence and MuSC activation, we performed a high throughput mRNA sequencing (mRNA-seq) experiment on 2 biological replicates of truly quiescent MuSCs isolated from male and female mice following *in situ* fixation (24). Principal component analysis revealed that samples clustered by genotype (PC 1) and by sex (PC 2) revealing potential sex differences in the response to glucocorticoids as has been previously reported (Fig. 4A) (46). When examining both males and females, the top differentially expressed genes, ranked by FDR using k means and k=6 revealed numerous cell cycle regulators (Fig. 4B). We next identified differentially expressed genes in females and males using a fold change cut-off of 1.5 and a false discovery rate (FDR) of less than 0.05 (Fig. 4C). In females, we identified 166 differentially expressed genes (97 upregulated, 69 downregulated) and in males, 196 differentially expressed genes were found (137 upregulated and 59 downregulated) (Fig. 4C). Interestingly, while both males and female mice display the same phenotype with loss of GR, we found only 8 significantly upregulated genes in common between males and females, including *Ccna2* and *Mki67*, two genes involved in cell proliferation, and 6 commonly downregulated genes including *Nr3c1* (Fig. 4C), suggesting that MuSC from males and females use unique pathways to cell cycle activation. The downregulation of *Nr3c1* (GR) in GR^MuSC-/-^ mice as compared to WT controls confirms the validity of the differential expression analysis.

**Figure 4.**
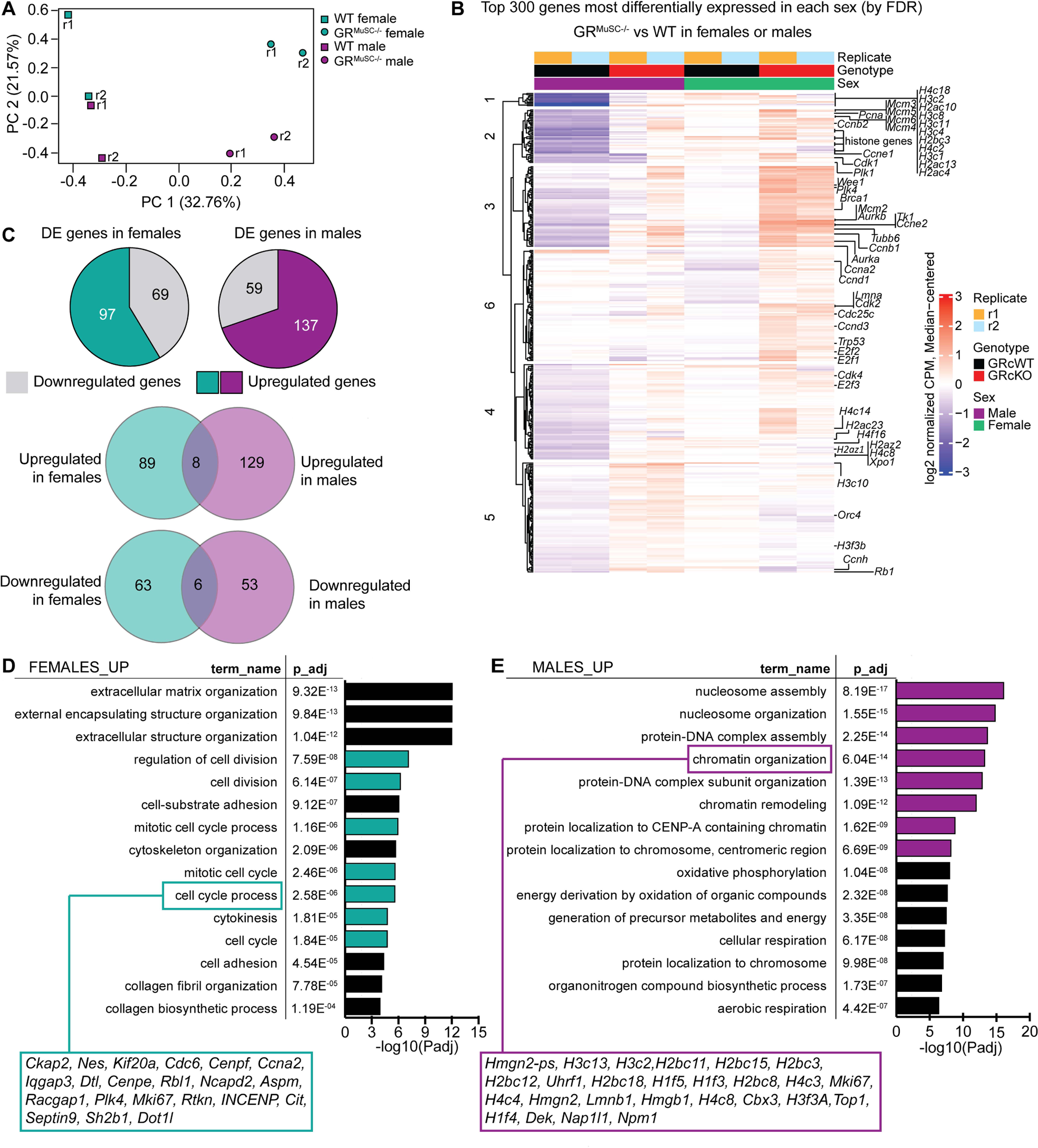
RNA-seq analysis of *in situ* fixed MuSCs reveals sex-differences with loss of GR expression. **(A)** Principal Component Analysis (PCA) plot for RNA-seq samples derived from *in situ* fixed MuSCs isolated from female and male WT and GR^MuSC-/-^ mice, n=2 samples sequenced, each derived from 6 WT or 6-7 GR^MuSC-/-^ mice. **(B)** A median-centered log2 normalized expression heatmap of the top 300 genes most differentially expressed (by FDR) in either female or male mice. The position of genes known to participate in the cell cycle is indicated. The genes (rows) were clustered by k-means with k=6 and the members of each cluster were subjected to hierarchical clustering. **(C)** Total number of differentially expressed genes using a fold-change cut-off of 2 and an FDR cut-off of 0.05 from B. Intersection of upregulated and downregulated differentially expressed genes from male and female mice is shown. **(D)** Top gene ontology terms (biological processes) in genes significantly upregulated in female *in situ* fixed MuSCs compared to WT controls. Select genes from the term “cell cycle process” are highlighted. **(E)** Top gene ontology terms (biological processes) in genes significantly upregulated in male *in situ* fixed MuSCs compared to WT controls. Genes from the “chromatin organization” term are highlighted.

The top enriched GOBP terms in the upregulated gene list in GR^MuSC-/-^ females highlighted extracellular matrix organization and cell division-related processes (Fig. 4D). In particular, the “cell cycle process” term, which included 21 genes, was enriched for genes that regulate the transition to M phase and cytokinesis. In males, the top enriched GOBP terms related to chromatin organization and metabolic processes, in particular, oxidative phosphorylation (Fig. 4E). Interestingly, the 26 genes included in the term “chromatin organization” were enriched for replication-dependent histones, histones that are expressed in a cell cycle specific pattern and support DNA replication during S phase.

Gene set enrichment analysis (GSEA) of the differentially expressed genes in female and male GR^MuSC-/-^ cells versus WT revealed significant enrichment of gene sets related to cell cycle regulation, including the REACTOME cell cycle core gene set in both females and males (Fig. 5A). Notably, only a few gene sets were differentially enriched in a sex-specific way. For example, the gene sets related to the gene ontology terms “Microtubule Cytoskeleton Organization” and “Regulation of Organelle Organization” were enriched in females, whereas the REACTOME gene sets “DNA Replication”, “Rho GTPase effectors”, “S phase” and the Hallmark gene set “Myc targets” were enriched in males (Fig. 5A,B and Supplemental Figure 4), consistent with our gene ontology analysis of differentially expressed genes in each sex. While sex differences were observed in our dataset, both males and females induce a gene expression pattern that is consistent with cycling cells suggesting that in the absence of GR, MuSCs exit quiescence.

**Figure 5.**
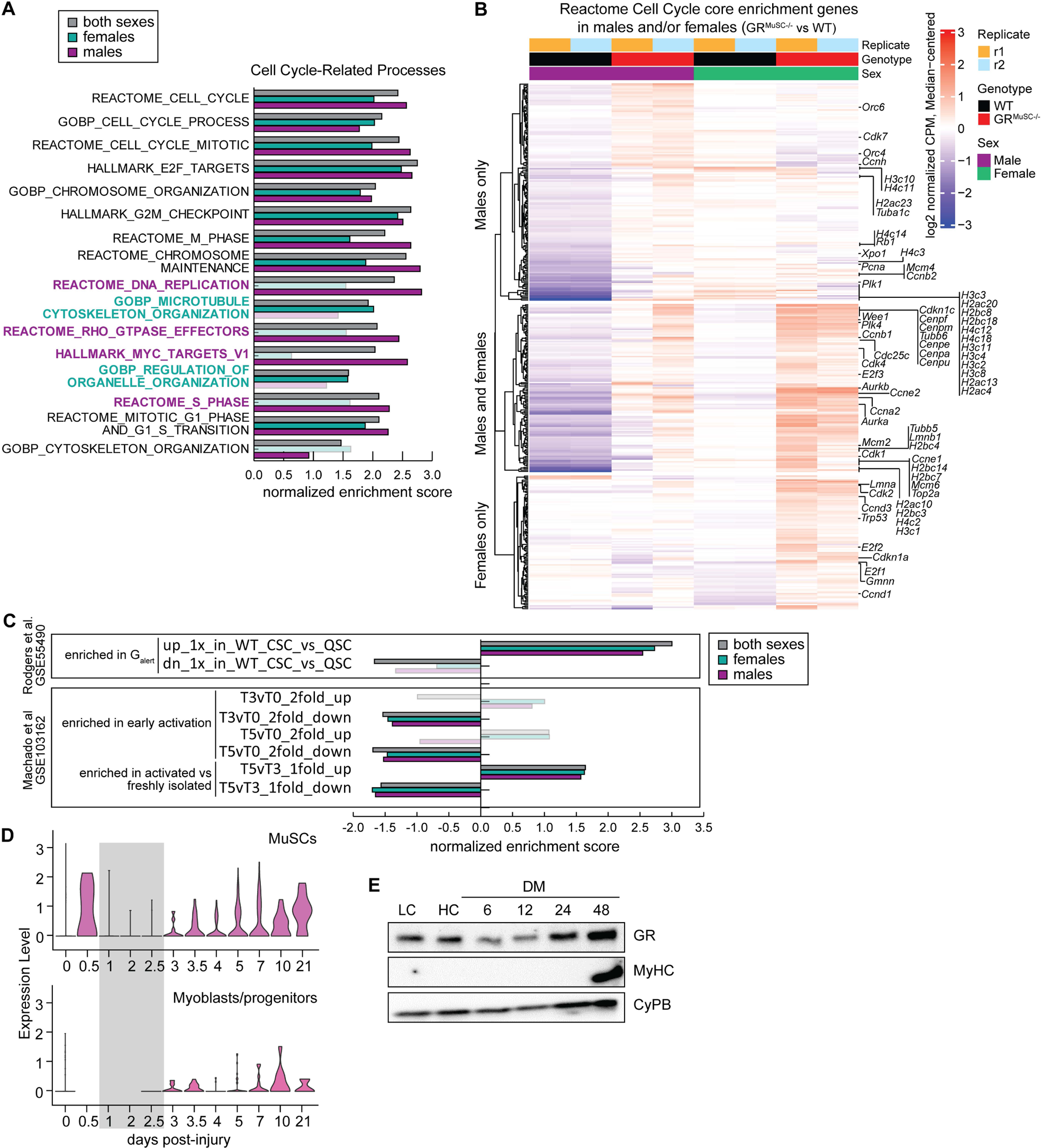
*In situ* fixed MuSCs from GR^MuSC-/-^ mice are precociously activated. **(A)** Gene set enrichment analysis (GSEA) of RNA-seq datasets from *in situ* fixed WT and GR^MuSC-/-^ MuSCs for females, males, and both sexes. Normalized enrichment scores with a p-adjusted value of >0.05 are transparent. Gene sets with sex-specific enrichment are highlighted. **(B)** Heatmap of enriched genes in female and/or male GR^MuSC-/-^ versus WT MuSCs using the REACTOME cell cycle gene set. **(C)** GSEA of RNA-seq data from females, males and both sexes compared to gene sets derived from Rodgers et al (GSE55490) and Machado et al (GSE103162). Transparent bars fail to meet statistical significance (P-adj >0.05). **(D)** *Nr3c1* mRNA expression during muscle regeneration in MuSCs and myoblast/progenitors using the scMuscle single cell transcriptomics database (48). **(E)** Western blot of GR and myosin heavy chain (MyHC) expression in growing MuSC-derived primary myoblasts at low confluency (LC) or high confluency (HC) or following induction to differentiate in differentiation medium (DM) for the hours indicated. n=3 biological pairs.

### GR^MuSC-/-^ MuSCs are more activated than WT MuSCs in the absence of injury

MuSCs in the muscles contralateral to injured muscle (contralateral satellite cells, CSCs) partially activate in response to the distant injury and enter a state known as G_alert_, which is distinct from MuSCs in injury-naïve mice (quiescent MuSCs, QSCs) and from injured tissue (activated MuSC, ASCs) (GSE55490) (47). We performed GSEA to compare the gene signature of *in situ* fixed GR^MuSC-/-^ with that of genes that are differentially expressed in CSC MuSCs (G_alert_) versus QSC MuSCs (Fig. 5C). We found significant positive enrichment in our dataset with genes that are upregulated in G_alert_ in both males and females (Fig. 5C). We also noted significant negative enrichment scores for downregulated genes in G_alert_ in both sexes (Fig. 5C and Supplemental Figure 5).

To evaluate the degree of activation in MuSCs lacking GR, we compared our lists of differentially expressed genes to the Machado *et al* dataset (Figure 1, GSE103164) (24). GSEA revealed significant negative enrichment scores for genes downregulated with early activation (T3) versus quiescence (T0), in essence, genes whose expression is associated with quiescence (Fig. 5C). Similar observations were made for GSEA of genes differentially regulated in growing MuSCs (T5) compared to quiescence, suggesting that GR positively regulates genes associated with the quiescent state rather than inhibit expression of activation genes. When our dataset was compared to genes differentially regulated between T5 and T3, we observed significant positive enrichment scores for both sexes for genes that are upregulated in T5 versus T3 (Fig. 5C). Further, we see significant negative enrichment scores for genes downregulated under these conditions (Fig. 5C). Interestingly, *Nr3c1* expression drops precipitously in MuSC and myoblasts early in repair after acute injury (48) (Fig. 5D) and GR protein expression is reduced during myogenic differentiation in culture (Fig. 5E) suggesting that in normal activation of MuSCs, they become temporarily insensitive to GCs. Thus, our findings support that the GR positively regulates genes that promote quiescence and is unlikely to suppress the expression of genes associated with activation. Further, our findings support that GR^-/-^ MuSCs are more similar to growing MuSCs than freshly isolated cells and have a gene signature that is consistent with the G_alert_ state.

### MuSC lacking the GR are more activated/cycling

To explore if MuSCs lacking the GR can return to quiescence, we performed immunostaining for Ki67 in injury-naïve muscle (Fig. 6A-C). In the absence of myotrauma, we did not expect to see many cycling cells and consistent with this, ∼10% of PAX7^+^ were Ki67^+^ in WT mice (Fig. 6B, C). In the uninjured GR^MuSC-/-^ muscle, we observed ∼21% positivity for Ki67, suggesting that GR^MuSC-/-^ cells are more activated than WT in injury-naïve animals (Fig. 6B, C). To confirm that MuSC lacking the GR are more proliferative, we injected 6-week-old mice for 5 consecutive days with tamoxifen to knock out GR expression and two weeks following the last injection, pulsed GR^MuSC-/-^ and WT mice with BrdU for 24h. Hindlimb muscles were collected a day later to assess efficiency of excision (Fig. 6D). Immunostaining for PAX7 and BrdU of TA muscle cross sections from WT and GR^MuSC-/-^ mice revealed that loss of the GR *in vivo* leads to a 3-fold increase in BrdU incorporation in females during the 24h pulse (Fig. 6E, F). This effect was not observed in male mice, suggesting potential differences in the timeline of activation. Taken together, these findings support a model in which the GR promotes the expression of genes that promote quiescence and, in its absence, precocious activation of MuSCs is observed.

**Figure 6.**
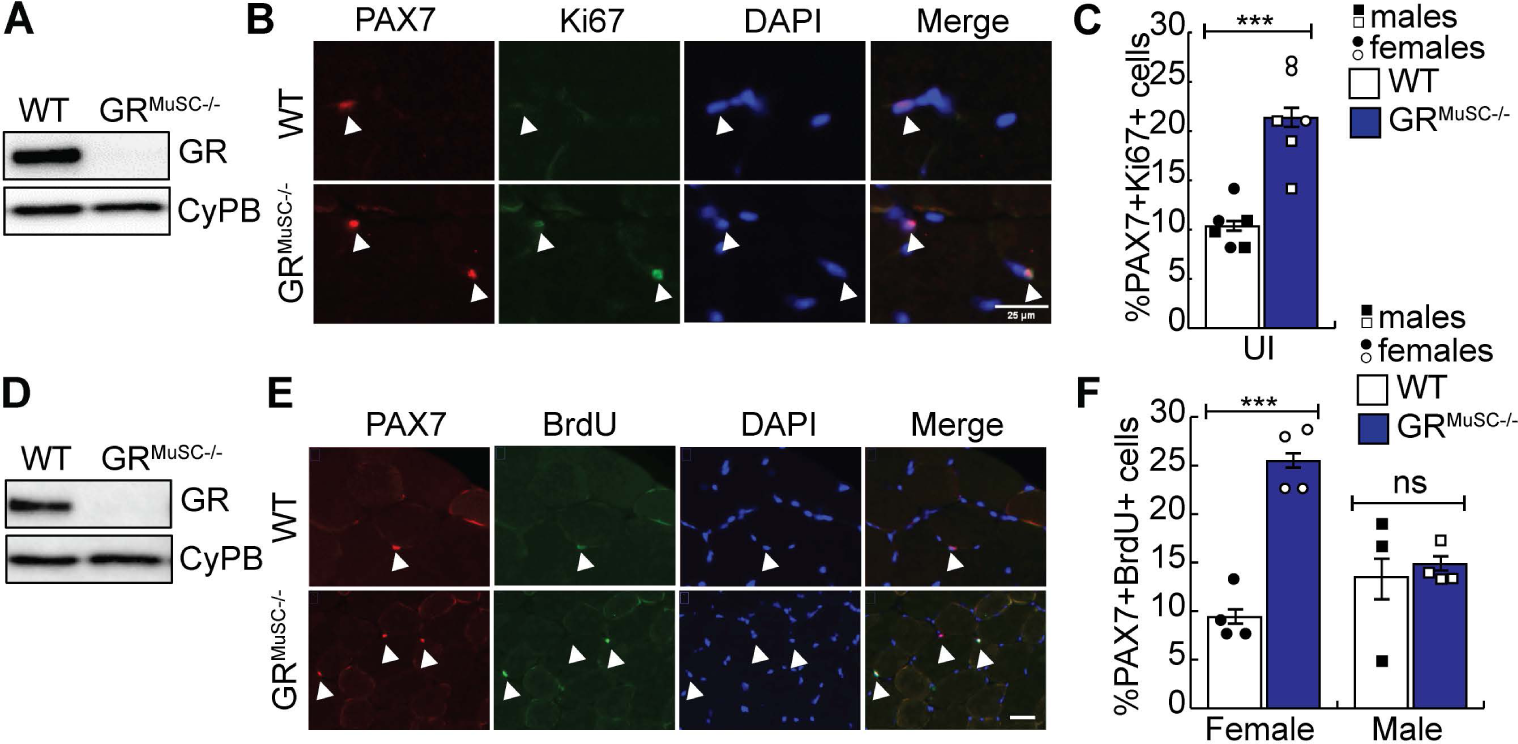
Loss of the GR increases MuSC cycling in the TA muscles of injury-naïve mice. **(A)** Western blot of GR expression in primary myoblasts isolated from uninjured GR^MuSC-/-^ and WT mouse 2 weeks after completion of tamoxifen administration. **(B)** Representative images of TA muscle sections stained for PAX7 (red) and Ki67 (green) from injury-naïve mice (UI). Nuclei were counterstained with DAPI (blue). Scale bar = 25 μm (UI). Arrows indicate the location of PAX7^+^ cells. **(C)** Percent PAX7^+^Ki67^+^ MuSCs from B, n=6 biological pairs. **(D)** Western blot of GR expression in primary myoblasts isolated from the GR^MuSC-/-^ and WT mouse 2 weeks after completion of tamoxifen administration and following a 24-hour pulse with BrdU prior to sacrifice. Cyclophilin B (CyPB) is a loading control. **(E)** Representative images of TA muscle sections stained for PAX7 (red) and BrdU (green) from D. Nuclei were counterstained with DAPI (blue). Scale bar = 25 μm. **(F)** Percent PAX7^+^BrdU^+^ cells from D, n= 4 biological pairs. Data is represented as mean ± SEM, ***p <0.001, two-tailed.

To identify GR targets genes in MuSC with a role in quiescence, we analyzed DNA accessibility regions from our ATAC-seq dataset to identify differentially regulated regions in quiescence versus early activation (Fig. 7A). From the 105010 peaks with above background ATAC signal, we identified 15,402 peaks that were differentially accessible using a logFC cutoff of >2 and a FDR of <0.05. We found that the majority of peaks (73%) had decreased accessibility with activation and these mapped to 1606 glucocorticoid response elements (GREs) near 1804 genes. Gene ontology analysis revealed enrichment for GOBP terms related to cell proliferation including “negative regulation of cell population proliferation” and “muscle cell proliferation” (Fig. 7B). We next explored RNA expression of the genes included in the muscle cell proliferation GO term in GR^MuSC-/-^ *in situ* fixed MuSC as compared to controls and in the Machado dataset (Fig. 7C). Of the genes with the highest expression in quiescent cells (T0) in this gene set and whose expression is downregulated with activation, we found significant downregulation of *Pik3r1*, *Tgfbr3*, *Stat5b*, *Npr3*, *Tenm4* and *Tgfbr2* in male and female GR^MuSC-/-^ cells. Of these, *Pik3r1* is a known GR target and has been shown to contribute to GC-mediated insulin resistance in chronic GC treatment (49). Further, PIK3R1 acts as a negative regulator of p100 enzymatic activity, a protein required for exit from quiescence by MuSCs (50).

**Figure 7.**
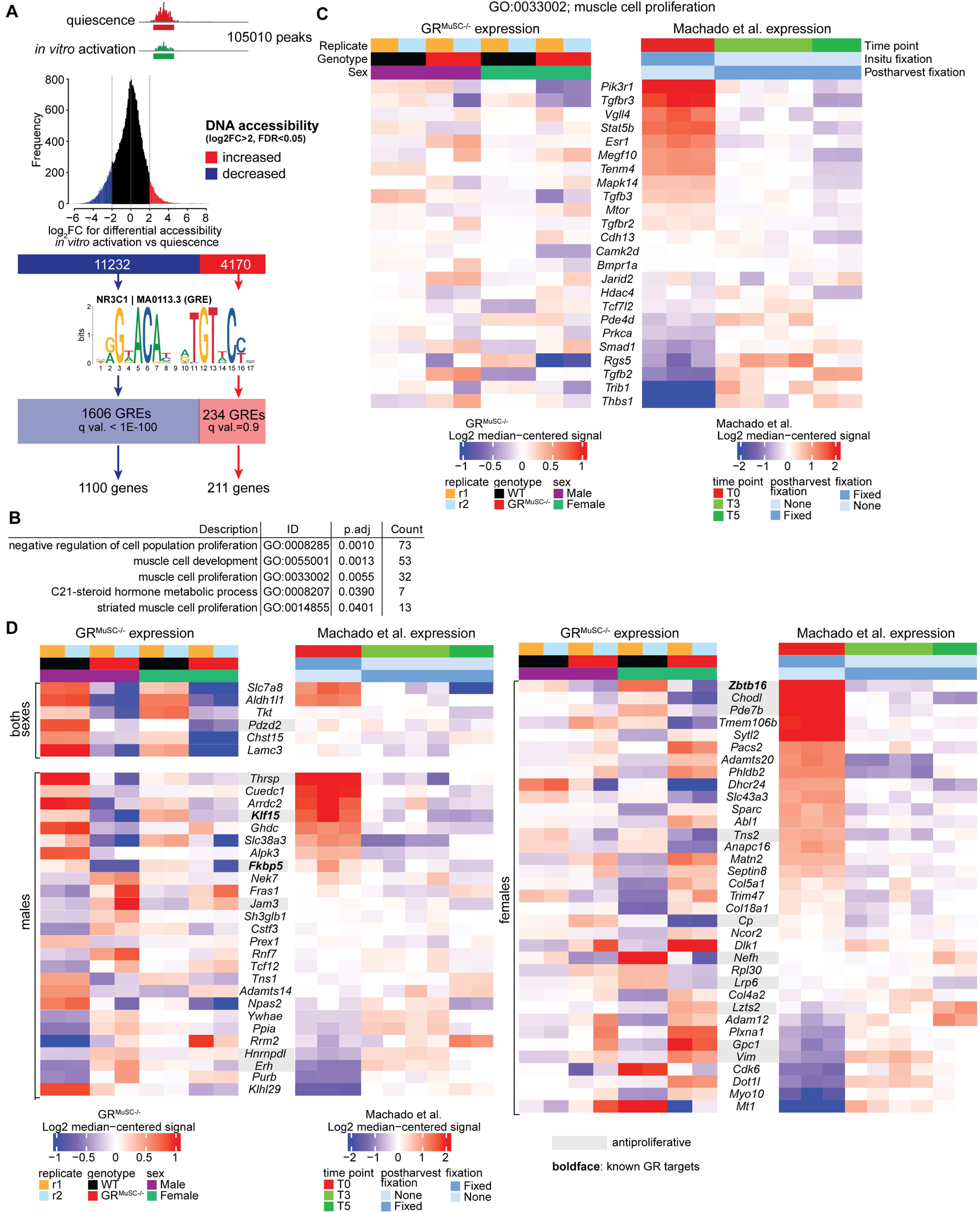
GR regulates genes that inhibit cell proliferation to promote quiescence. **(A)** Analysis workflow to identify GR target genes from ATAC-seq datasets in freshly isolated versus quiescent MuSCs using mapping of Glucocorticoid response elements (GREs). **(B)** Gene ontology (GOBP) analysis of putative GR target genes identified in a from the 1100 genes near GREs located in regions of maximum DNA accessibility in quiescent cells. The 211 genes near GREs located in regions with maximum DNA accessibility in activated cells were not enriched for any GOBP term, at the FDR < 0.05 cutoff. **(C)** RNA expression of genes identified as regulators of muscle cell proliferation in b in WT and GR^MuSC-/-^ MuSCs and in quiescent, freshly isolated and growing MuSCs from the Machado dataset. *Pik3r1* is a known GR target. **(D)** Genes bearing GREs identified in a were cross-referenced with differentially expressed genes in quiescent MuSCs from WT and GR^MuSC-/-^ mice using a cut-off FDR of 0.1 and compared to expression data from the Machado dataset. Genes in boldface are known GR targets. Genes highlighted in grey have a described anti-proliferative role.

Next, we compared genes with reduced DNA accessibility with activation, generated using a cut-off of log2 FC>2 and FDR<0.05, to differentially expressed genes in males and female GR^MuSC-/-^ cells using a FDR cut-off of 0.1 (Fig. 7D). This comparison revealed several genes that are most highly expressed in quiescent cells in the Machado dataset (T0) and downregulated in the GR^-/-^ MuSCs including *Slc7a8*, *Aldh1l1*, *Thrsp*, *Klf15*, *Fkbp5*, *Zbtb16*, *Chodl*, and *Pde7b* among others (Fig. 7D). Globally, this analysis reveals that the gene expression pattern of identified GR targets in both sexes is consistent with an activation gene signature and reveals novel GR targets that regulate MuSC quiescence.

## Discussion

Our findings place GR as a critical regulator of MuSC quiescence and suggests that the primary function of GR in this capacity is to stimulate the expression of anti-proliferative genes rather than inhibit proliferation genes. For example, in both sexes, loss of GR increases *Pdzd2* expression, which has been shown to act as a negative regulator of proliferation (51). *Zbtb16*, a known GR target that mediates GC-induced cell cycle arrest (52, 53) and *Chodl*, a negative regulator of proliferation that is preferentially expressed in muscle (54, 55) are significantly downregulated in female GR^-/-^ MuSCs however, the pattern of expression is conserved in males. In males *Fkbp5* and *Klf15*, two known GR targets (56, 57) have the same expression profile. FKBP5 inhibits Cdk4 and thereby the cell cycle to promote myogenic differentiation and has been shown to be anti-proliferative in multiple models (58–61). *Klf15* is also a negative regulator of proliferation (62, 63) and has been shown to regulate myotube size downstream of GCs (64), however, a role for this factor in the regulation of MuSC function has not been described.

Another interesting GR target revealed in our analysis is *Pik3r1. Pik3r1* has been identified as a target in myoblasts (65) and overexpression of *Pik3r1* in myotubes reduces cell diameter (49). Knockdown of *Pik3r1* in myotubes impairs GC-mediated suppression of insulin signalling (49). Functionally, *Pik3r1* encodes a negative regulator of PI3K, acting to block the enzymatic activity of the p110 subunit (66), which, in turn, has been shown to be both necessary and sufficient for quiescence exit in adult MuSC (50). Consistent with the phenotype of GR*^MuSC-/-^* mice, constitutive PI3K signalling results in precocious exit from quiescence and nuclear accretion via fusion to existing myofibers in the absence of injury (50, 67, 68). Conditional knockout of P110α in MuSCs prevents MuSCs from exiting quiescence following injury and impairs muscle regeneration (50, 68). PI3K activity inhibits FOXO3A activity and stimulates mTORC, which has been implicated in the regulation of the G_alert_ state (67). Thus, our findings support a model in which GR is required to promote the expression of *Pik3r1* which inhibits PI3K activity to promote MuSC quiescence. Following injury or loss of GR, *Pik3r1* expression is downregulated, resulting in enhanced PI3K activity, and ultimately the exit from quiescence.

In a broader sense, it is interesting to intersect the role of GCs in the stress response in the context of myotrauma and their role in the regulation of quiescence in MuSCs. In multiple models of injury, endogenous GCs have been shown to be rapidly and transiently secreted (69, 70) and this action is vital for the coordination of the body’s response to trauma, with low cortisol levels predictive of poor outcomes (71). Our findings suggest that rising GC levels following injury would prevent MuSC activation in the early response to myotrauma. While the impact of GC release following trauma on MuSCs is unknown, it may serve to protect this vital stem cell population from activation in a muscle microenvironment that cannot support their differentiation. Another intriguing possibility is that the transient high levels of GCs following injury serve to coordinate MuSC proliferative responses to injury by synchronizing the tissue-level circadian clock. Indeed, cell cycle progression is regulated by circadian rhythms (72, 73) and the circadian rhythm regulator *Per3*, which is proposed to have tissue-level timekeeping effects (74), is significantly upregulated with loss of GR in MuSCs in both males and females. Intestinal stem cells have been shown to proliferate in response to injury in a circadian and Per-dependent manner (73, 75), suggesting that GCs may act to synchronize transcriptional responses in early regeneration and during activation. Indeed, GCs have been implicated in the regulation of peripheral, tissue-level clocks in lung (76). Consistent with this, we find that the GR is downregulated in early regeneration and myogenic differentiation, suggesting that decreased sensitivity to GCs contributes directly to MuSC activation following injury.

In conditions with chronically high levels of GCs due to pharmacological treatment or illness, we predict stem cell defects, characterized by precocious differentiation, impaired MuSC self-renewal, reductions in stem cell populations and eventually impaired regenerative responses. Indeed, exposure to GCs during fetal development results in smaller MuSC populations in rats and consequently, impaired postnatal growth of skeletal muscle fibers (77). Further, the synthetic GC dexamethasone was shown to inhibit muscle regeneration in pigs, resulting in fewer myogenic precursor cells one day post-injury (78). While not specifically investigated in this study, the depletion of MuSC observed in the GR^MuSC-/-^ mouse following injury suggests that GR may be required for both the maintenance and the establishment of quiescence, such that in its absence, self-renewal is impaired. While GR expression is transiently downregulated in differentiating MuSCs, it is unclear if the receptor becomes re-expressed in self-renewing cells and the impact of GC levels on this process. In a severe mouse model of Duchenne Muscular Dystrophy (DMD), GC treatment reduced MuSC depletion, suggesting that GCs may promote self-renewal and quiescence. The therapeutic benefits of GC in DMD may thus be attributed to beneficial effects on the myofiber and its resilience (79), which result in less regenerative pressure, as well as protection of the stem cell population.

While our findings indicate that MuSCs from both male and female mice precociously activate in response to loss of GR expression, the transcriptomes are not identical. The small number of commonly differentially regulated genes suggest that the response to GCs is sexually dimorphic. In support of this, distinct transcriptional responses to GCs have been described in muscle and other systems (46, 80, 81). GR has been shown to overlap with estrogen receptor-mediated responses and can interfere with them, providing a potential molecular mechanism for the sex-based differences in transcriptional responses observed (82, 83). In muscle, GC treatment resulted in improved muscle performance in both males and females but achieved this effect by stimulating different pathways including enhanced lipid metabolism in females that improved endurance over males (46). While published literature focuses on the effects of GCs on the myofiber, our work indicates that distinct transcriptional responses exist in the MuSC population. Given that GCs are the primary treatment for X-linked congenital muscular dystrophies such as Duchenne Muscular Dystrophy, it is imperative to fully understand the implications of GC treatment on the MuSC population in the context of these myopathies. For example, GC therapy has been shown to be beneficial for approximately 3 years after which muscle wasting accelerates (84). A close examination of MuSC function in the context of this therapeutic window may provide clarity as to the mechanism of treatment failure, as our results would support poor proliferative responses and abnormal self-renewal in the context of GCs that would negatively impact long-term muscle regeneration.

## Data Availability

### Funding

This work was supported by the Canadian Institutes of Health Research (CIHR) [PJT 175164 to NWB, PJT 183839 to AB] and the Muscular Dystrophy Association [Idea grant #874651 to AB and NWB]. RR is supported by the Ontario Graduate Scholarship (OGS) from the University of Ottawa, ON. AS is supported by a Queen Elizabeth II Graduate Scholarship in Science and Technology.

## Acknowledgements

The authors wish to thank Leanne Dawe, Dabo Yang and Jocelyn Nguyen for assistance with sample collection, Philippos Mourikis (Institut Mondor, Paris) for advice on RNA extraction from in situ-fixed samples, the uOttawa Cell Biology and Image Acquisition Core (RRID: SCR_021845) for help with image acquisition, processing, and quantification, the Louise Pelletier Histology Core (RRID: SCR_021737) for H&E staining on TA muscles used in this study, and the uOttawa Flow Cytometry Core Facility for assistance with MuSC isolation. This research was enabled in part by support provided by WestGrid and the Digital Research Alliance of Canada (<u>alliancecan.ca</u>).

## Author Contributions

RR performed the majority of the experiments and performed the analysis (investigation, analysis and validation). RR participated in the drafting of the original manuscript and data visualization. HA assisted in data collection and analysis, aided in the conceptualization of the project, drafting of the original manuscript and data validation. AS assisted in data collection, performed data analysis and reviewed the manuscript. AB assisted in data collection, performed data analysis and reviewed the manuscript. IB assisted in the development of methodology (investigation) and reviewed the manuscript. AB* participated in the conceptualization of the project, the supervision of the project, data visualization, data acquisition and analysis, data curation and drafting of the manuscript. AB* also provided research funding for this project. NWB was primarily responsible for the conceptualization of the project, supervision, funding, data visualization and writing of the manuscript.

