## Supplemental Figures for "The glucocorticoid receptor is a critical regulator of muscle satellite cell quiescence"

**Rajgara *et al.* Supplemental Information**

**Supplemental Figure Legends**

**Supplemental Figure 1. Profiling of DNA accessibility in MuSCs that are quiescent or subjected to *in vitro* activation. (A)** Heatmap of sample-to-sample distances, calculated using Pearson correlation coefficients. Rows and columns represent cell samples subjected to ATAC-seq. "Quiescent" refers to cells subjected to *in situ* fixation, "Fresh" to cells allowed to undergo activation during the isolation process, and "Growing" to cells put in culture in growth medium for 24 hours prior to the ATAC procedure. **(B)** Principal component analysis showing sample clustering based on ATAC-seq signal at all peaks discovered. Samples cluster primarily by whether they underwent activation or not, and post-harvest fixation has only a limited effect (diamonds are close to circles of the same color). **(C)** Example of ATAC-seq signal for genes that have regions with either higher (right-hand side) or lower (left-hand side) DNA accessibility after *in vitro* activation. The RNA-seq data from Machado et al. is also shown for reference. The RNA-seq is presented in a strand-specific manner, with read density on the plus strand shown in green, and on the negative strand shown in purple. The "early activation" time point represents T3 in Machado et al.

**Supplemental Figure 2. DNA accessibility and gene expression profiles in quiescent and early-activation MuSCs.** Related to Figure 1a. Boxplots are used to show the log2 fold-change in DNA accessibility (from ATAC-seq) or mRNA levels (from RNA-seq) between fresh (early activation) and in situ-fixed (quiescent) cells for the regions and genes represented in each of the twelve clusters. The number of differentially accessible regions represented in each cluster is indicated in blue.

**Supplemental Figure 3. Gene set enrichment analysis (GSEA) of RNA-seq samples derived from in situ fixed WT and GR$^{MuSC-/-}$ MuSCs.** Enrichments that fail to meet statistical significance are transparent. Enriched gene sets with sex differences are highlighted.

**Supplemental Figure 4. Differentially expressed genes in MuSCs lacking GR are enriched for cell cycle genes. (A)** A median-centered log2 normalized expression heatmap of the top 300 genes most differentially expressed in *in situ* fixed MuSCs from RNA sequenced from WT and GR$^{MuSC-/-}$, ranked by FDR without imposed cut-off. n=2 samples derived from multiple pooled female mice. **(B)** Running enrichment score (ES) plot of all differentially expressed genes in female *in situ* fixed MuSC (contrast GR$^{MuSC-/-}$ vs WT compared to the REACTOME "Cell Cycle" gene set of 610 genes). **(C)** A median-centered log2 normalized expression heatmap of the top 300 genes most differentially expressed in *in situ* fixed MuSCs from RNA sequenced from WT and GR$^{MuSC-/-}$, ranked by FDR without imposed cut-off. n=2 samples derived from multiple pooled male mice. **(D)** Running enrichment score (ES) plot of all differentially expressed genes in male *in situ* fixed MuSC (contrast GR$^{MuSC-/-}$ vs WT compared to the REACTOME "Cell Cycle" gene set of 610 genes).

**Supplemental Figure 5. Differentially expressed genes in MuSCs lacking GR are enriched for the Galert gene signature. (A)** Heatmap of the median-centered log2 normalized expression of the core genes upregulated in the Galert dataset (GSE55490) and enriched in female GR$^{MuSC-/-}$

*in situ* fixed MuSC versus WT. **(B)** Running ES plot using contrast $GR^{MuSC-/-}$ vs WT in female mice compared to genes upregulated in Galert. **(C)** Heatmap of core genes upregulated in the Galert dataset (GSE55490) enriched in male $GR^{MuSC-/-}$ mice versus WT. **(D)** Running ES plot using contrast $GR^{MuSC-/-}$ vs WT in male mice compared to genes upregulated in Galert.

**Supplemental Figure 6. Differentially expressed genes in *in situ* fixed MuSCs lacking GR are enriched for genes expressed in activated MuSCs.** **(A)** Heatmap of the median-centered log2 normalized expression of the core gene set from genes downregulated in freshly isolated female-derived MuSC versus in situ fixed (quiescence genes) (GSE103164, Machado T3 vs T0 RNA-seq data)). **(B)** Running ES plot of genes in females (contrast $GR^{MuSC-/-}$ vs WT) compared to downregulated quiescence genes from (A). **(C)** A heatmap of the median-centered log2 normalized expression of the core gene set from genes downregulated in freshly isolated male-derived MuSC versus in situ fixed (quiescence genes) (GSE103164, Machado T3 vs T0 RNA-seq data)). **(D)** Running ES plot of genes in males (contrast $GR^{MuSC-/-}$ vs WT) compared to downregulated quiescence genes from (A).

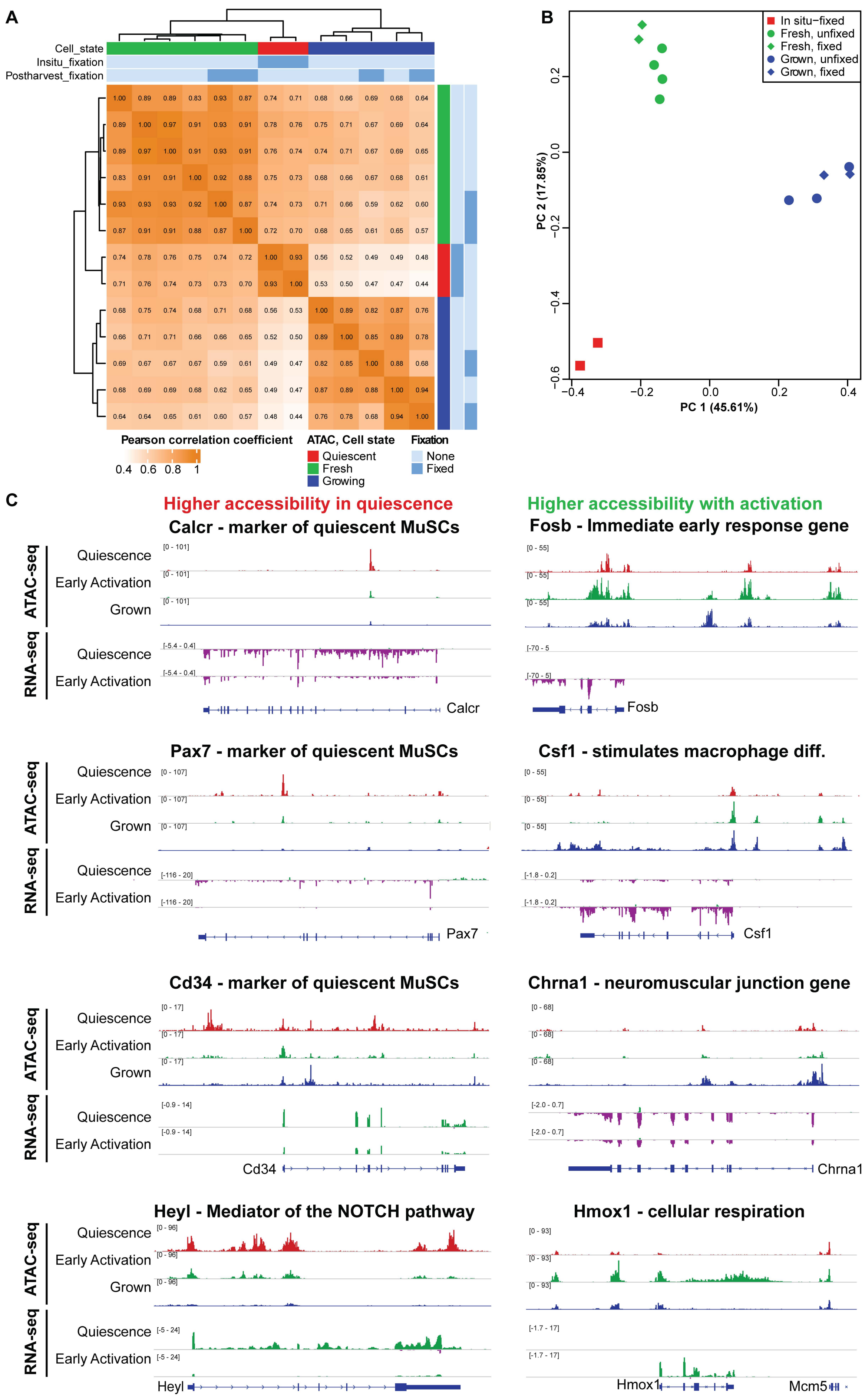


Rajgara et al. Supplemental Figure 1

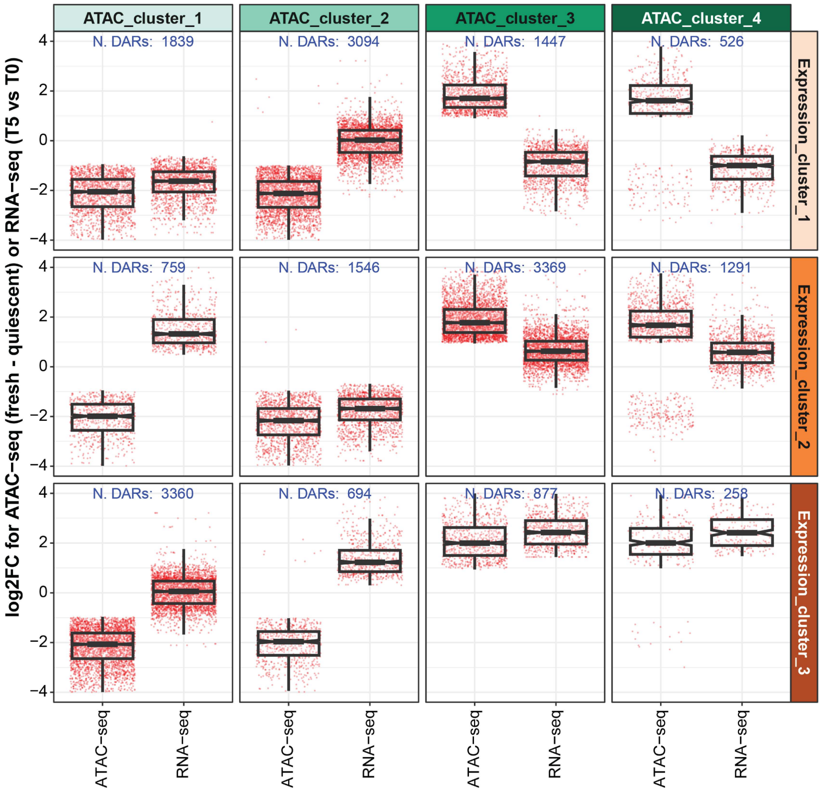


Rajgara et al. Supplemental Figure 2

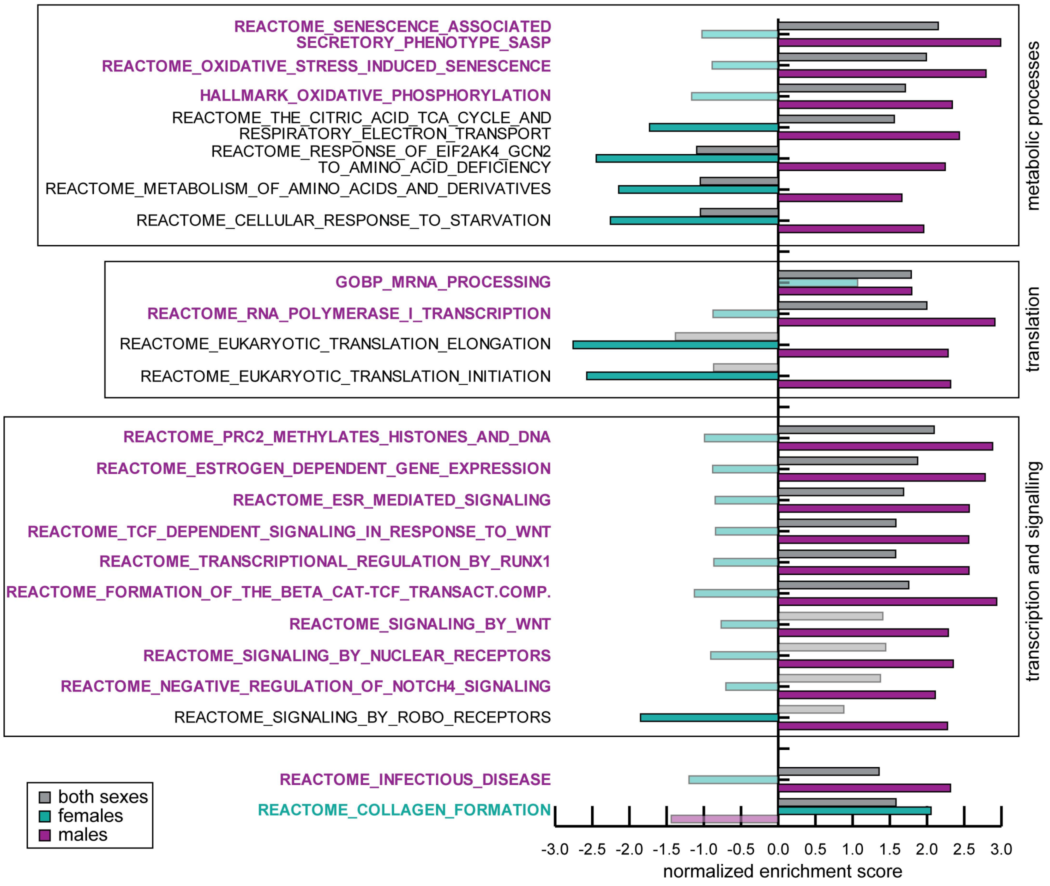


**Rajgara et al. Supplemental Figure 3**

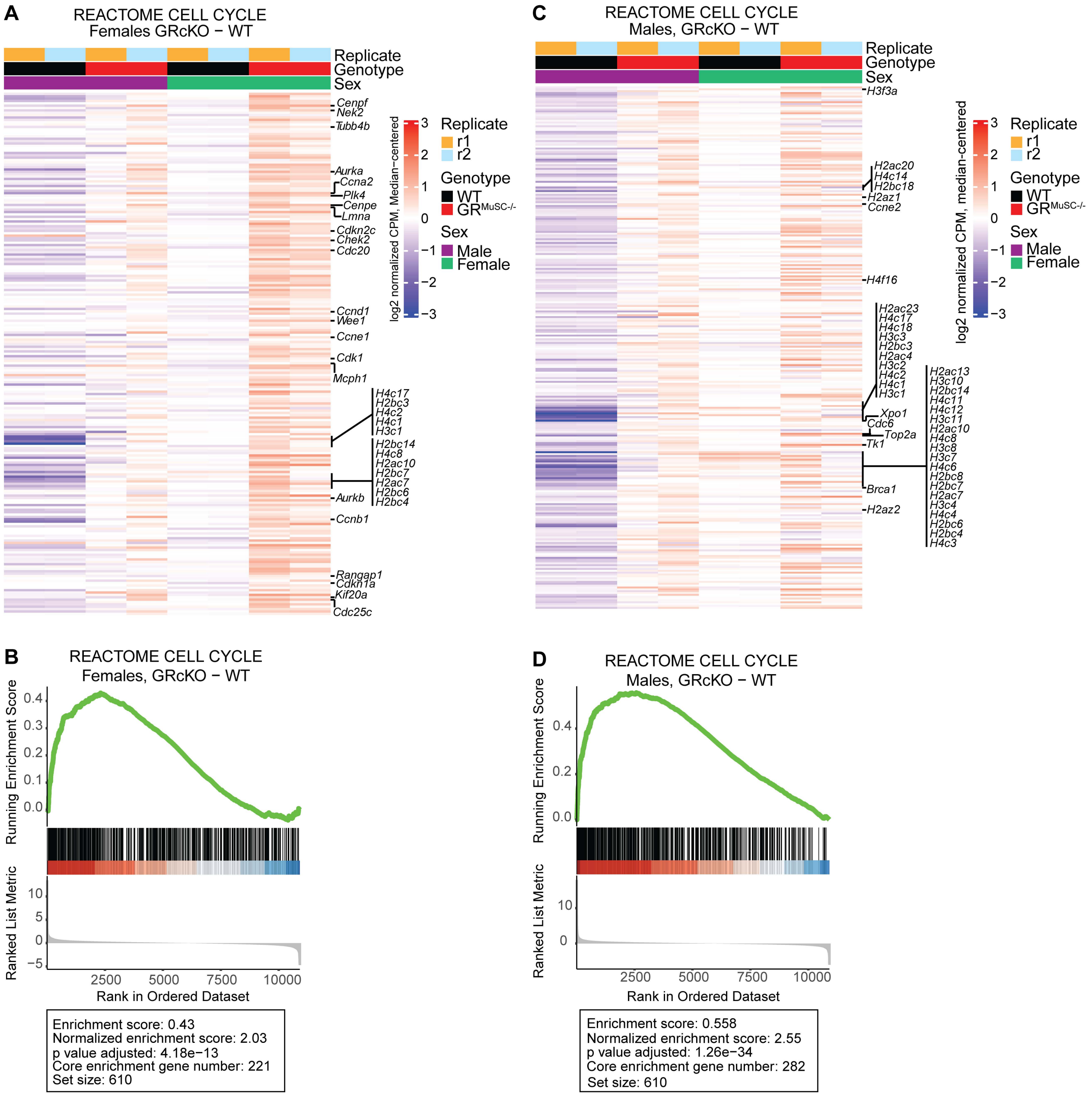


**Rajgara et al. Supplemental Figure 4**

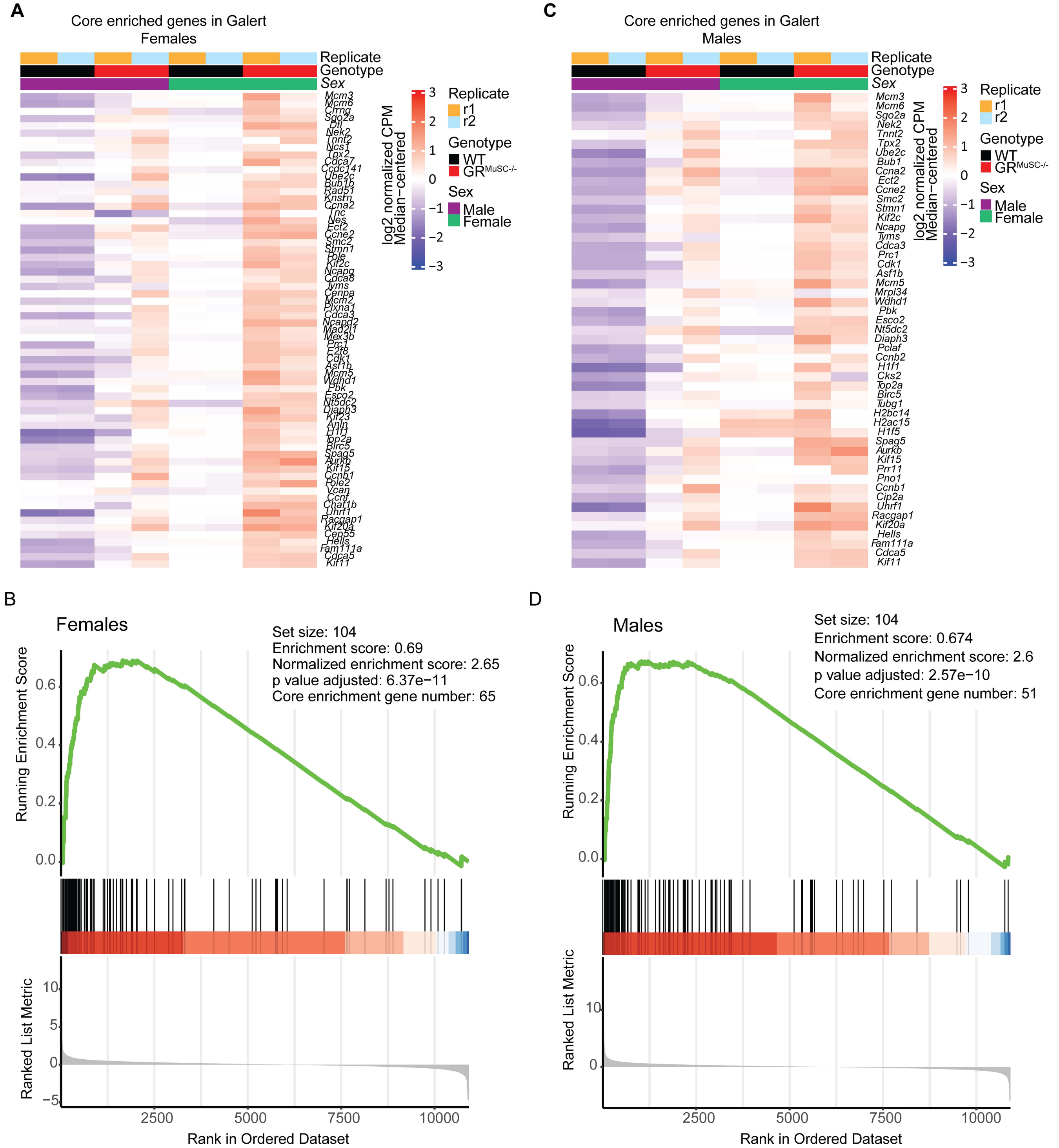


**Rajgara et al. Supplemental Figure 5**

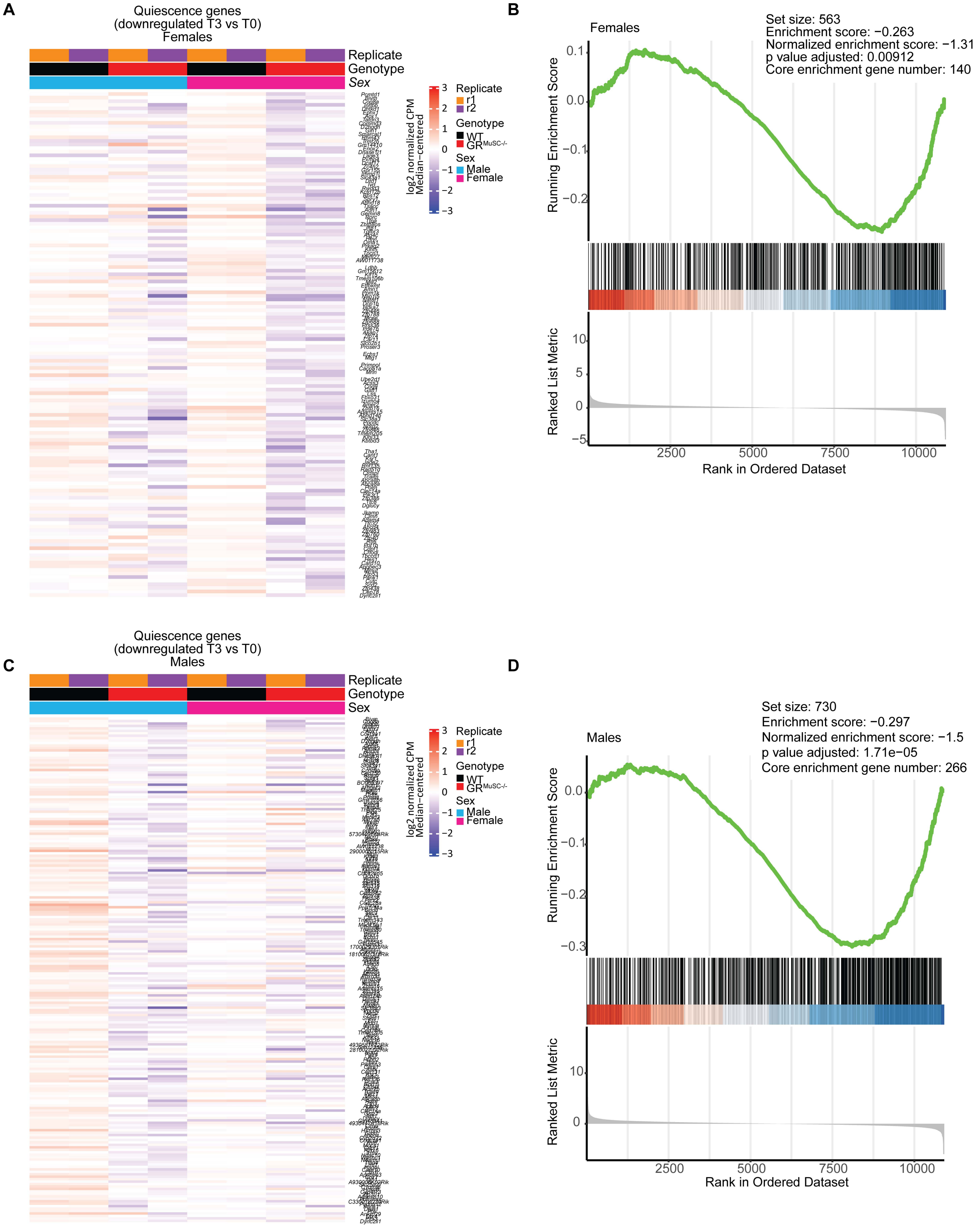


**Rajgara et al. Supplemental Figure 6**
